# Target-specific co-transmission of acetylcholine and GABA from a subset of cortical VIP^+^ interneurons

**DOI:** 10.1101/469064

**Authors:** Adam J. Granger, Wengang Wang, Keiramarie Robertson, Mahmoud El-Rifai, Andrea Zanello, Karina Bistrong, Arpiar Saunders, Brian Chow, Vicente Nuñez, Chenghua Gu, Bernardo L. Sabatini

## Abstract

The modulation of cortex by acetylcholine (ACh) is typically thought to originate from long-range projections arising in the basal forebrain. However, a subset of VIP interneurons express ChAT, the synthetic enzyme for ACh, and are a potential local source of cortical ACh. Which neurotransmitters these VIP/ChAT interneurons (VCINs) release is unclear, and which post-synaptic cell types these transmitters target is not known. Using quantitative molecular analysis of VCIN pre-synaptic terminals, we show expression of the molecular machinery to release both ACh and GABA, with ACh release restricted to a subset of boutons. A systematic survey of potential post-synaptic cell types shows that VCINs release GABA primarily onto other inhibitory interneuron subtypes, while ACh neurotransmission is notably sparse, with most ACh release onto layer 1 interneurons and other VCINs. Therefore, VCINs are an alternative source of cortical ACh signaling that supplement GABA-mediated disinhibition with highly targeted excitation through ACh.

## Introduction

In the mammalian cortex, acetylcholine (ACh) is a major neuromodulator that promotes attention and learning (Hasselmo and Sarter, 2011; Klinkenberg et al., 2011). High tonic levels of ACh correlate with arousal and alertness (Teles-Grilo Ruivo et al., 2017), whereas phasic increases in ACh coincide with the detection of important sensory cues (Parikh et al., 2007; Sarter et al., 2009, 2014). The net effect of activating cholinergic fibers in cortex is the desynchronization of neuronal firing (Pinto et al., 2013), and ACh can gate the induction of synaptic plasticity (Morishita et al., 2010; Rasmusson, 2000), enabling learning. In addition, ACh dilates cortical blood vessels (Lecrux et al., 2017; Yamada et al., 2001) allowing for greater delivery of oxygen and nutrients to support increased metabolic demand. In summary, ACh is a vital signaling molecule that coordinates cortical activation and plasticity.

The source of cortical ACh is typically attributed to long-range projections from cholinergic neurons of the basal forebrain (Mesulam et al., 1983). However, a population of local cortical neurons exists across species that could provide an alternative source of ACh. Initially identified by immunolabelling for choline acetyltransferase (ChAT)(Eckenstein and Baughman, 1984; Eckenstein and Thoenen, 1983), the biosynthetic enzyme that produces ACh, their presence has been repeatedly confirmed by immunohistochemical and transcriptional analyses (Bhagwandin et al., 2006; Cauli et al., 1997; Consonni et al., 2009; Gonchar et al., 2008; Kosaka et al., 1988; Peters and Harriman, 1988; Porter et al., 1998; Schäfer et al., 1994; Weihe et al., 1996). Although not widely appreciated, both classic studies and modern databases of single cell transcriptomes (Saunders et al., 2018; Tasic et al., 2016; Zeisel et al., 2015) demonstrate that cortical ChAT^+^ neurons express vasoactive intestinal peptide (*Vip*), suggesting that they are a subclass of VIP interneurons.

Despite being defined by the expression of a protein for ACh synthesis, it is unclear what neurotransmitters these neurons release. At least one study concluded that that they lack expression of the vesicular ACh transporter (VAChT) (Bhagwandin et al., 2006) and therefore are unable to release ACh. Conversely, given the expression of VIP and their potential identity as VIP interneurons, one would expect them to be GABAergic. Indeed, they have been shown to contain GABA (Bayraktar et al., 1997). However, other studies report GABAergic protein expression in only a subset of cortical ChAT^+^ neurons (von Engelhardt et al., 2007; Kosaka et al., 1988), and a preliminary exploration of their functional connectivity concluded that they primarily release ACh onto pre-synaptic nicotinic ACh receptors (nAChRs) to promote the release of glutamate by other neurons (von Engelhardt et al., 2007). Therefore, whether these neurons are competent to release ACh and/or GABA remains unclear.

We recently established that co-transmission of GABA is a common feature of cholinergic neurons in the mouse forebrain (Granger et al., 2016; Saunders et al., 2015a). GABA and ACh have opposite effects on membrane voltage through ionotropic receptors, though metabotropic receptors may have net excitatory or inhibitory effects depending on the receptor subtype and subcellular localization, as well as complex biochemical effects through second messengers. Given their potential to release neurotransmitters with opposing ionotropic effects, the functional consequence of these cells on cortical circuits is unknown.

In this study, we demonstrate the co-transmission of ACh and GABA from cortical ChAT^+^ neurons, which we confirm are a subset of VIP interneurons. Contrary to previous studies (Bhagwandin et al., 2006; von Engelhardt et al., 2007; Kosaka et al., 1988), we find that VIP/ChAT interneurons (VCINs) express genes necessary to synthesize and release both ACh and GABA. High-throughput imaging analysis of proteins within VCIN pre-synaptic terminals demonstrate co-localization of ACh and GABA release machinery, with ACh release restricted to a subset of terminals. Screening of post-synaptic connectivity showed that these neurons sparsely target specific cells with ACh and GABA, consistent with previously studied long-range cholinergic projections (Saunders et al., 2015b). We therefore systematically surveyed candidate post-synaptic targets of GABA and ACh co-transmission, testing responses from different interneuron subtypes, pyramidal neurons, pre-synaptic receptors, non-neuronal targets like cortical arteries, and homotypic connections between VCINs themselves. We find that GABA release is robust onto other inhibitory interneuron subtypes, consistent with the general connectivity of VIP interneurons, and that ACh release is sparse, with the strongest responses elicited in layer 1 interneurons and VCINs themselves. Thus, VCINs are a specialized subclass of VIP interneurons that specifically excite other VCINs and layer 1 interneurons while inhibiting many classes of cortical GABAergic interneurons.

## Results

### Characterization of Cortical VIP/ChAT interneurons

To characterize the morphology and laminar distribution of cortical ChAT^+^ neurons, we crossed Chat^ires-Cre^ (Rossi et al., 2011) and Rosa26^lsl-tTomato^ (Ai14) (Madisen et al., 2010) mice to induce expression of tdTomato in all cholinergic neurons (Figure 1A,B), and confirmed that the ChAT^ires-Cre^ line faithfully reports cortical *ChAT* expression (Figure S1). TdTomato-expressing cells were present in all cortical regions but clustered mostly in superficial layers (Fig. 1D). Most of the ChAT^+^ neurons have bipolar morphology (67%, Figure 1C,D), with dendrites extending vertically towards the pia and deep layers. A smaller subset have an irregular morphology, with 3 or more main dendrites extending radially and cell bodies that are concentrated at the border of layer 1 and 2/3 (33%, n = 102, Figure 1C,D). This morphology and laminar distribution is similar to that of VIP interneurons. Indeed, both immunohistochemistry and fluorescent *in situ* hybridization (FISH) show that nearly all cortical ChAT^+^ neurons express VIP, and cortical ChAT^+^ neurons comprise 30-50% of VIP interneurons (Figure 1E,F). We saw no coexpression with parvalbumin (PV) or somatostatin (Sst) (Figure S2), indicating that that ChAT^+^ neurons are an exclusive subset of VIP interneurons. We further confirmed that cortical ChAT^+^ interneurons are a subset of VIP interneurons using publically-available single-cell RNA sequencing of cortical neurons from the Allen Brain Institute (Figure S3; data from (Tasic et al., 2018), accessible at http://celltypes.brain-map.org/rnaseq/mouse). Therefore, we refer to these cells as VIP/ChAT interneurons (VCINs).

**Figure 1.**
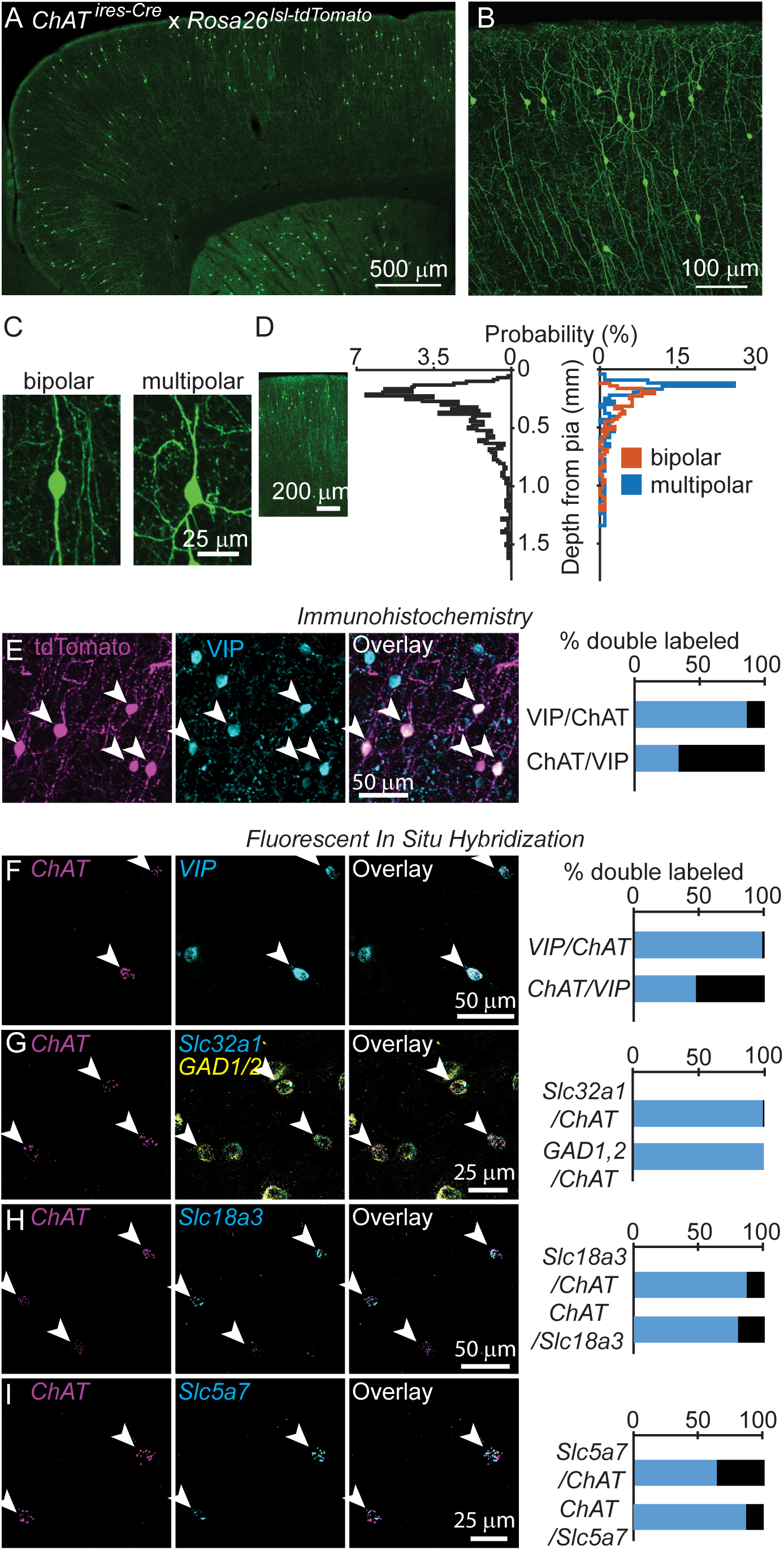
Cortical ChAT+ neurons are a subset of VIP interneurons that express GABAergic and Cholinergic genes. A,B) Example sagittal images of cortical ChAT+ neurons, labeled in ChAT^ires-Cre^x Rosa26^lsl-tdTomato^ mice demonstrating cellular distribution (A) and morphology (B). C) Example images of a bipolar (left) and multipolar (right) morphology). D) Distribution of cortical depth from pia of all cortical ChAT+ neurons (left graph, black trace, n = 1059 neurons, median cell body is 274 µm from pia ± 15 µm, 95% C.I.) and according to morphology (right graph; orange = bipolar, n = 207, 66% of total, median 293 µm from pia ± 23 µm, 95% C.I.; blue = multipolar, n = 107 neurons, 34% of total, median 173 µm from pia ± 24 µm, 95% C.I.). E) Example images (left) and quantification (right) of cortical ChAT+ interneurons from ChAT^ires-Cre^ x Rosa26^lsl-tdTomato^ mice (left panel) colabeling with immunostained VIP (middle panel), and the overlay (right panel; n = 127 ChAT+,VIP+ neurons of 147 total ChAT+ and 375 VIP+ neurons). F) Fluorescent in situ hybridization against ChAT in cortex (left panel) overlaps with VIP (middle panel; overlay, right panel; n = 278 ChAT+,VIP+ of 283 ChAT+ and 579 VIP+ neurons). G) ChAT expression in the cortex co-labels with the GABAergic genes Slc32a1, which encodes for VGAT, and a combined label for Gad1,Gad2, which encodes for GAD67,65 (n = 101 ChAT+,Slc32a1+ and 102 ChAT+,GAD1/2+ of 102 ChAT+ neurons). H,I) ChAT expression in the cortex co-labels with the Cholinergic genes Slc18a3, which encodes for VAChT, and Slc5a7, which encodes for ChT (n = 147 ChAT+,Slc18a3+ of 170 ChAT+ and 184 Slc18a3+ neurons; n = 72 ChAT+,Slc5a7+ of 113 ChAT+ and 83 Slc5a7+ neurons). Arrowheads indicate cortical ChAT+ neurons.

Whether all VCINs express the genes necessary for synthesis and release of both GABA and ACh is unclear. Single-cell RNA sequencing data indicates that all VIP interneurons express GABAergic genes (Figure S3) and labeling of VCINs by FISH shows they all express mRNAs encoding the vesicular GABA transporter (VGAT, encoded by *Slc32a1*), and GABA synthetic enzymes GAD65/67 (encoded by *Gad1/2*, Figure 1G). Although VCINs by definition express the ACh synthetic enzyme ChAT, they may not express the remaining cellular machinery necessary for ACh release, such as the membrane choline transporter (ChT, encoded by *Slc5a7*) and vesicular ACh transporter (VAChT, encoded by *Slc18a3*). Again, both FISH and single-cell RNA sequencing data indicate that *ChAT+* neurons of the cortex also express *Slc18a3* and *Slc5a7* (Figure 1H,I, Figure S3). These data show that VCINs express the genes necessary for synthesis and synaptic release of both GABA and ACh.

### Molecular characterization of VCIN pre-synaptic terminals

Somatic expression of mRNAs encoding ACh and GABA transmission machinery neither guarantees that VCINs express these proteins at pre-synaptic release sites nor informs what types of synapses they make. To characterize the proteins present at the pre-synaptic terminals of VCINs, we used array tomography (Micheva and Smith, 2007), which permits multi-plexed immunolabeling of many synaptic epitopes. To label pre-synaptic terminals, we injected AAV(8)-DIO-Synaptophysin-YFP into the frontal cortex of *ChAT^ires-Cre^* mice (Saunders et al., 2015b)(Figure 2A,B), and analyzed expression of 7 synaptic proteins in addition to using DAPI to mark cell nuclei and Synaptophysin-YFP fluorescence to identify the pre-synaptic terminals of VCINs. Specifically, we used Synapsin-1 as a general pre-synaptic marker, PSD-95 and VGLUT1 to label glutamatergic synapses, Gephyrin and VGAT to label GABAergic synapses, and ChAT and VAChT to label cholinergic synapses (Figure 2B,C).

**Figure 2.**
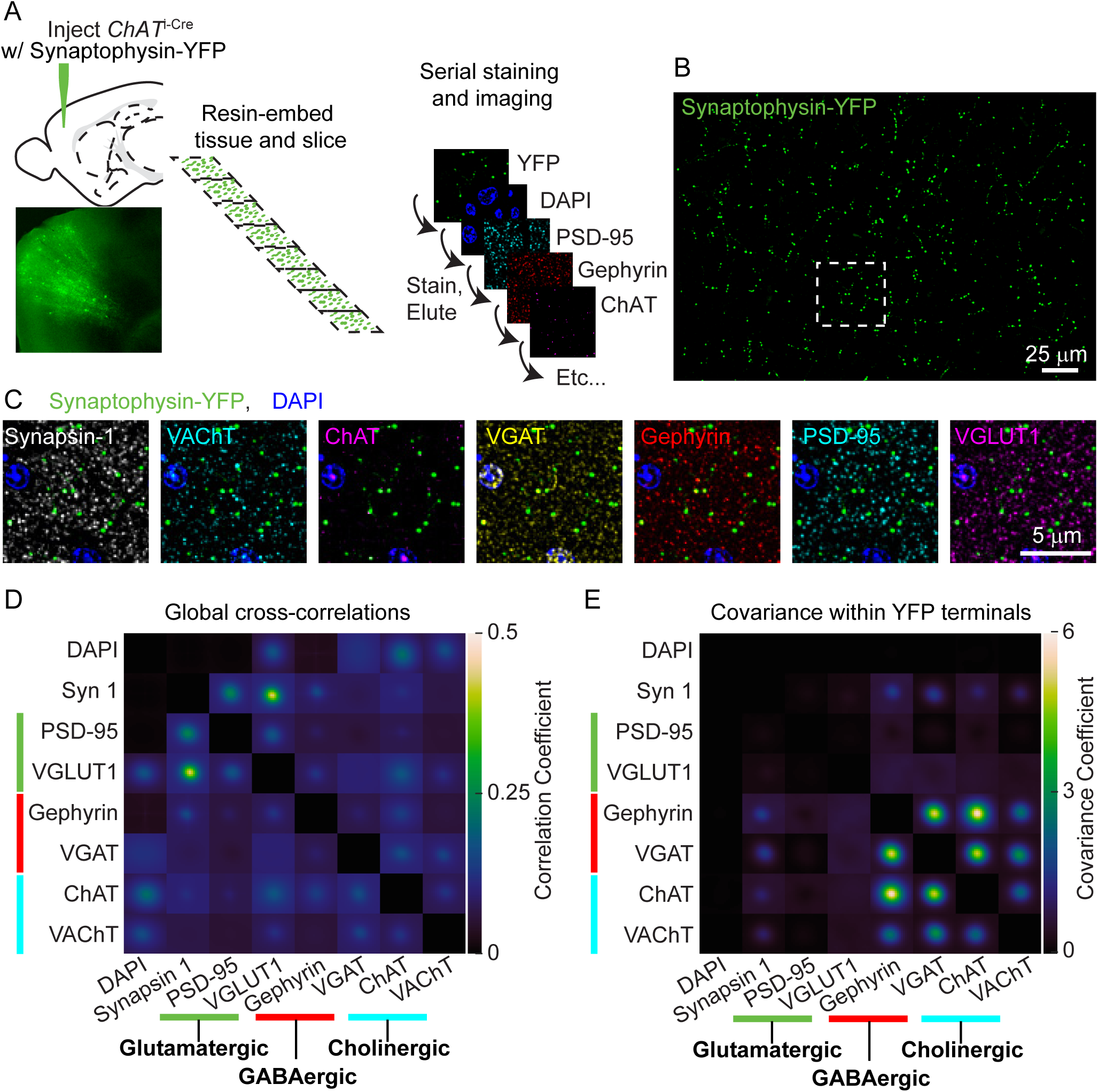
Multiplexed labeling of synaptic proteins within the pre-synaptic terminals of VCINs shows correlations between GABAergic and Cholinergic proteins. A) Array tomography workflow: ChATires-Cre mice were injected with AAV(8)-DIO-Synaptophysin-YFP to label the pre-synaptic terminals of VCINs. Approximately 1 mm^2^ squares of tissue were embedded in resin and cut into ultra-thin, 70 nm slices in an array. The ribbons of tissue were serially stained with antibodies against pre- and post-synaptic proteins. B) Example maximum projection of the Synaptophysin-YFP staining. C) Inset from (B) showing the staining of individual pre- and post-synaptic proteins. D) The average cross-correlation of across all pairs of raw images of pre- and post-synaptic antibody stains (n = 8 image stacks from 3 mice). E) The average co-variance across all pairs of pre- and post-synaptic antibody stains only within a mask created by the Synaptophysin-YFP signal. Co-variance within YFP terminals is not limited to values between -1 and 1 because all antibody stains were z-scored prior to masking and calculating co-variance, and therefore antibody signals may be more or less concentrated within the YFP mask (n = 8 image stacks from 3 mice).

To test whether specific pairs of synaptic markers have similar patterns of expression, as well as to determine the level of non-synaptic antibody staining, we calculated the global cross-correlations across all pairs of image arrays of fluorescent antibodies (Figure 2D, also see (Micheva and Smith, 2007)). Any correlation with DAPI, which labels cell nuclei where no synapses are present, is indicative of non-synaptic labeling. In general the antibodies for synaptic markers avoided cell nuclei, although VGLUT1, VGAT, ChAT, and VAChT did show small positive correlations with DAPI (Figure 2D). The strongest cross-correlations occurred between Synapsin-1, PSD-95, and VGLUT1, which is an indication both of the high density of excitatory synapses and the relatively low background signal with these antibodies. However, when considering lower density cholinergic and GABAergic synapses, the background signal obscures potential significant cross-correlations in the entire image arrays. In striking contrast, when we restrict our analysis to the ~0.1% area of the images covered by VCIN-expressed Synaptophysin-YFP (see Methods), we see high covariance among the GABAergic and cholinergic markers, with little to no covariance with the glutamatergic markers (Figure 2E). Thus fluorescence of pre-synaptic markers of ACh and GABA release are correlated within the pre-synaptic terminals of VCINs, indicating that individual terminals have machinery to release both ACh and GABA, but as expected for VIP neurons, not glutamate. In addition, the high covariance of Gephyrin specifically with VAChT and ChAT shows that VCIN terminals typically form inhibitory synapses.

While GABAergic and cholinergic markers co-vary specifically within VCIN terminals, these relationships do not inform whether GABAergic and cholinergic proteins are enriched in VCIN terminals compared to the rest of the cortex. To rigorously quantify enrichment of these synaptic proteins in VCIN terminals, we used the YFP signal to define binary 3D masks corresponding to the volumes of putative pre-synaptic terminals. We assigned each punctum of antibody signal to an individual pixel corresponding to the location of its peak fluorescence and quantified the chance that an antibody punctum colocalized with each voxel within the pre-synaptic terminals (Figure 3A). To determine if an antibody is enriched in terminals, we compared its percent colocalization within the YFP^+^ terminals to that within the volume around each terminal. In addition, we compared the actual colocalization with the distributions of chance colocalization measured by randomizing the locations of the antibody puncta 1000 times (Figure 3A). For example, for a single sample, Synapsin 1 is significantly enriched within the pre-synaptic terminals above chance levels, but not in the area surrounding the pre-synaptic terminals (Figure 3B). In contrast, PSD-95 and VGLUT1 are colocalized with YFP^+^ mask significantly below chance levels, while both GABAergic and cholinergic markers were found strongly above chance (Figure 3C). Across all samples, Synapsin-1, Gephyrin, VGAT, ChAT, and VAChT were consistently enriched within VCIN terminals, while PSD-95 and VGLUT1 were specifically depleted (Figure 3D). Because both VAChT and VGAT expression are central to our conclusions, we validated these two antibodies with genetically mosaic conditional knockouts and found that both their high covariance and enrichment within VCIN terminals were eliminated when we selectively deleted VGAT and VAChT in VCINs (Figure S4 and S5).

**Figure 3.**
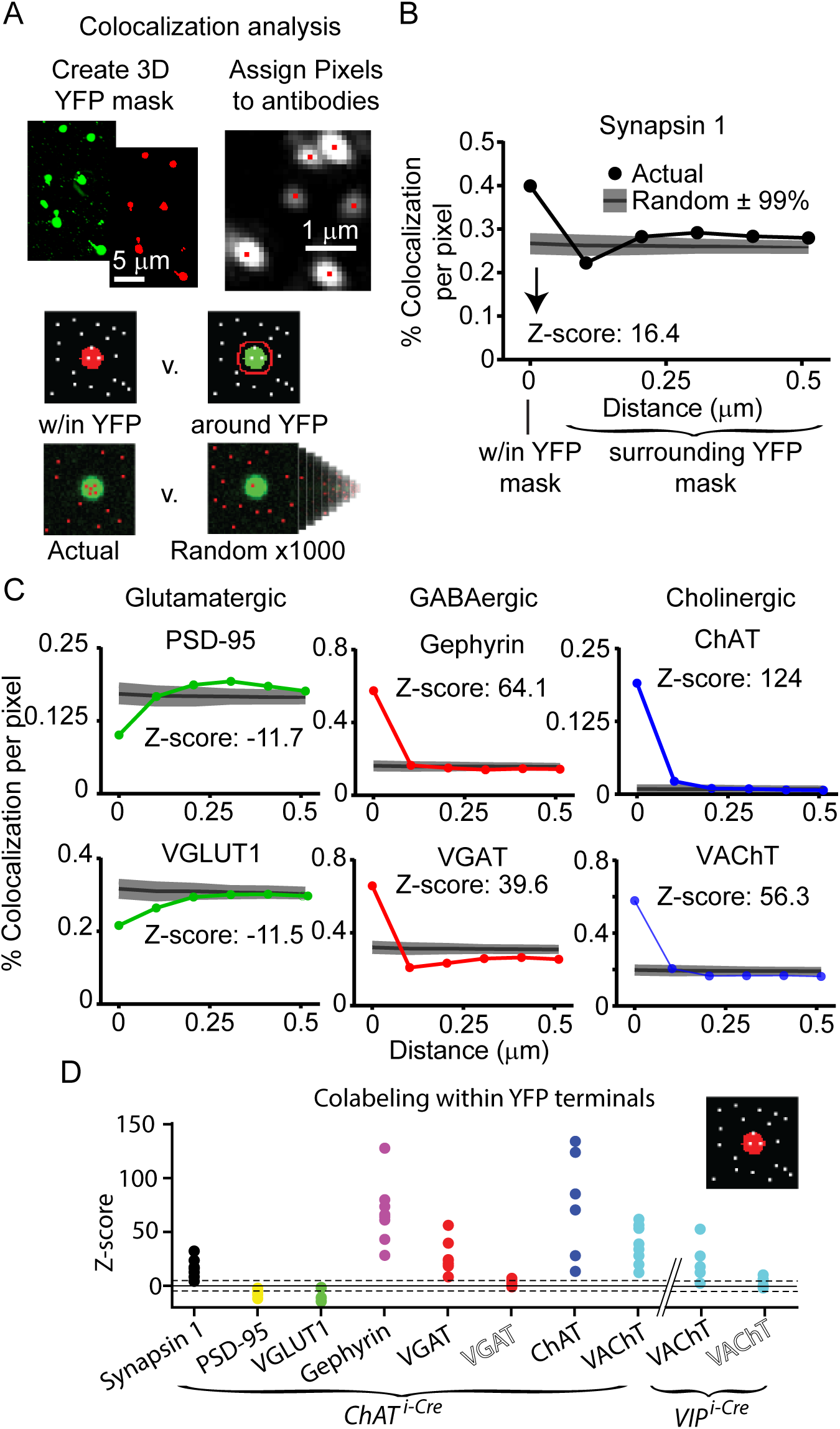
GABAergic and Cholinergic synaptic proteins are enriched within the pre-synaptic terminals of VCINs. A) Analysis workflow: YFP-expressing pre-synaptic terminals are converted into 3D mask volumes and each antibody puncta is assigned a pixel location based on the location of peak fluorescence. The percentage of voxels that overlap with an identified antibody pixel are quantified within the YFP terminals and in the volume around the terminals. These colocalization percentages are compared with the colocalization from 1000 rounds of randomized antibody puncta locations. B) Example graph showing the actual percent colocalization of Synapsin 1 puncta with the YFP mask, and in expanding 0.1 µm volumes around the EYFP mask. Actual colocalization is overlaid with the mean ± 99% of 1000 rounds of randomized puncta locations, which is used to calculate a z-score of antibody enrichment above or below chance. C) Examples from a single sample showing the colocalization of glutamatergic, GABAergic, and cholinergic markers with EYFP-expressing pre-synaptic terminals of VCINs. D) Colocalization z-scores across antibodies for all samples. Tissue samples from ChATires-Cre mice are shown, as well as the VGAT antibody z-score from VGAT conditional knock-out mice (indicated by VGAT in outline lettering, from ChAT^ires-Cre^ x VGAT^fl/fl^ mice), and VAChT antibody z-scores from VIP^ires-Cre^ and VIP^ires-Cre^ x VAChT^fl/fl^ (indicated by VAChT in outline lettering). Dashed lines indicate ± 5 z-scores.

The array tomography data shows that VCINs broadly express GABAergic and cholinergic proteins within their terminals and associate with inhibitory synapses, but do not indicate whether there is heterogeneity among terminals regarding which neurotransmitters are released. We therefore turned to traditional three-color IHC to classify the terminals as GABAergic (by expression of VGAT) or cholinergic (by expression of VAChT). Traditional IHC shows highly specific labeling against VGAT and VAChT (Figure 4A), and the thicker slices compared to array tomography allow for easier tracking of individual axons with many putative pre-synaptic terminals. This analysis revealed that pre-synaptic terminals varied considerably in their expression of VAChT (Figure 4B-D). Typically, an entire axon branch either expresses VAChT or not (Figure 4B,C). However, we also observed individual axons with intermingled pre-synaptic terminals that were either VAChT^+^ or VAChT^-^ (Figure 4D). We quantified the mean fluorescent intensity of VGAT and VAChT within each mCherrry-expressing terminal, and observed a large proportion of terminals had very low VAChT fluorescence intensity, overlapping with negative control intensities calculated by rotating the pre-synaptic terminal ROIs with respect to the VAChT signal (Figure 4F). We also observed a correlation between the intensity of VGAT and VAChT across terminals (R-squared = 0.33), although there appears to be two populations of terminals, those in which VAChT intensity scales with increased VGAT fluorescence and those in which VAChT intensity does not increase with increased VGAT (Figure 4E). To systematically classify each terminal as positive or negative for each marker, we set a fluorescence intensity cut-off that best differentiates VAChT or VGAT signal from background for each image. Using this classification, the majority of putative terminals were judged to express both VAChT and VGAT (Figure 4G, R-squared = 0.232). However, a large subset expressed VGAT and not VAChT (Figure 4G, R-squared = 0.091). We confirmed that this relationship held true across a range of classification thresholds (Figure S6A,B). As a negative control, we repeated this analysis in Synaptophysin-mCherry expressing terminals of Sst^+^ interneurons and observed no VAChT^+^ terminals, and no relationship between VGAT and VAChT fluorescence intensity per bouton (Figure S6D-H). In summary, we conclude that VAChT is present in only a subset of pre-synaptic terminals of VCINs, and therefore the release of ACh from these neurons is likely to be targeted to a specific population of post-synaptic cells.

**Figure 4.**
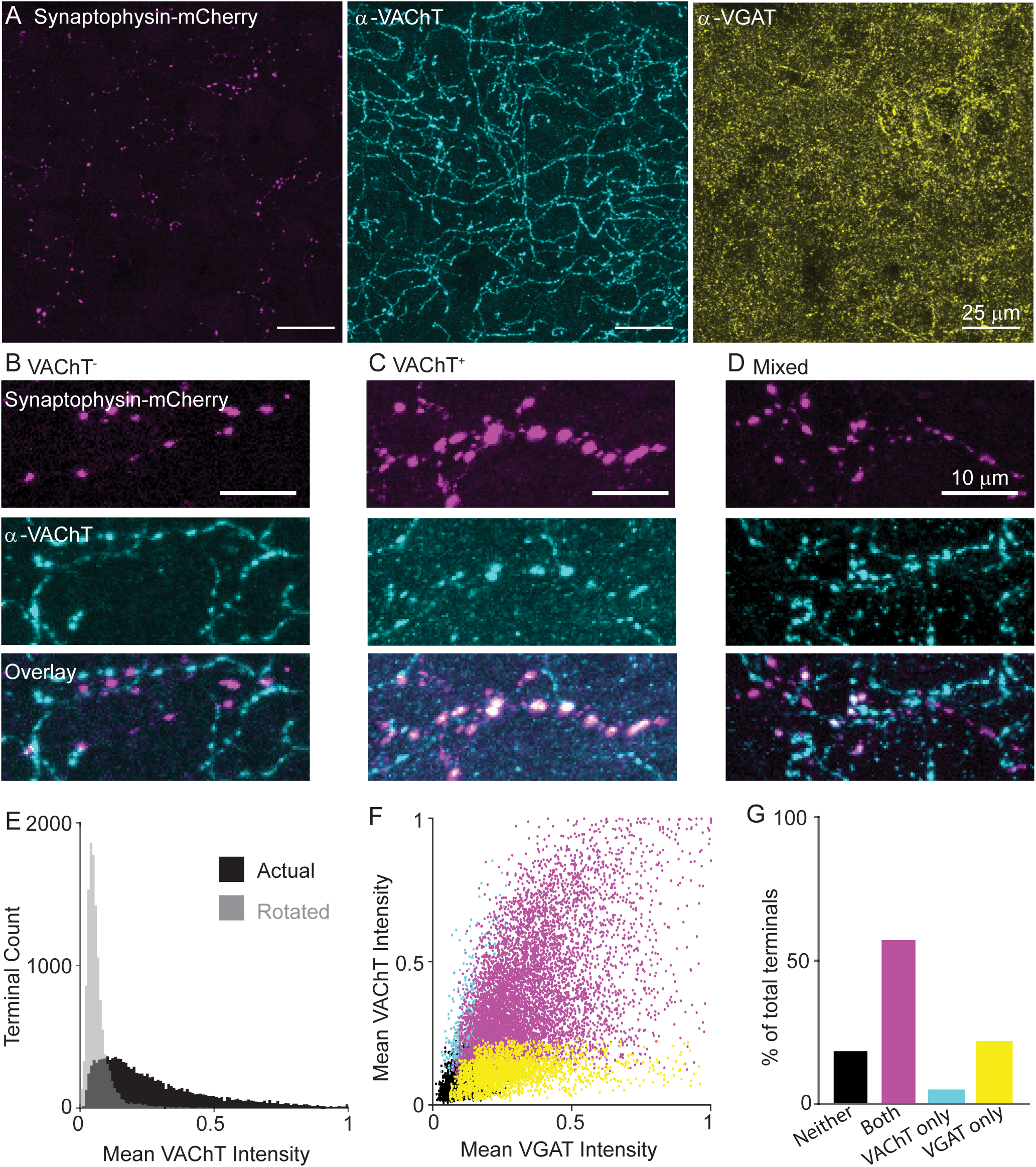
VCINs express VAChT in a subset of pre-synaptic terminals. A) Example images of VCIN pre-synaptic terminals labeled with AAV(8)-DIO-Synaptophysin-mCherry injected in the frontal cortex of ChATires-Cre mice. Left: Synaptophysin mCherry; Middle: VAChT immunostain; Right: VGAT immunostain. B-D) Example images showing putative Synaptophysin-mCherry+ terminals, showing largely VAChT- terminals (A), VAChT+ terminals (B), and a intermingled terminals that are both VAChT+ and VAChT-. (C). E) Histogram of mean VAChT fluorescence intensity within Synaptophysin-mCherry+ terminals. Black histogram represent the actual VAChT intensities, grey histogram represents the mean VAChT intensities when the mCherry mask is rotated 90° relative to the VAChT immunostain. F) Scatter plot of mean VGAT intensity and VAChT intensity in each putative pre-synaptic terminals (n = 12,356 putative terminals from 30 image stacks from 3 mice). Terminals are color-coded according to expression of VAChT and VGAT (Black – neither VGAT or VAChT, Magenta – both VGAT and VAChT, Cyan – VAChT only, Yellow – VGAT only). G) Quantification of the number of terminals of each type in (F).

### Post-synaptic targets of VIP/ChAT interneurons

The molecular analyses indicate that VCINs likely co-transmit GABA and ACh with segregated release of ACh from only some of their axon terminals. However, it is unknown what the specific post-synaptic targets of GABA and ACh co-transmission are. To determine the post-synaptic connectivity of VCINs, we injected AAV(8)-DIO-ChR2-mCherry into the frontal cortex of *ChAT^ires-Cre^* mice to enable selective optogenetic activation of VCINs. After 3-weeks of ChR2 expression, we obtained whole-cell recordings in acute brain slices from ChR2-lacking neurons and determined if exciting VCINs with blue light evoked inhibitory or excitatory post-synaptic responses (Figure 5A). GABA- and ACh-mediated ionotropic responses were differentiated by holding the postsynaptic neuron at 0 mV and -70 mV, respectively. In several post-synaptic neurons we observed excitatory post-synaptic currents (EPSCs) at -70 mV, which were confirmed to be mediated by nicotinic ACh receptors (nAChRs) by their unique kinetics (Figure 2B, see (Bennett et al., 2012) and sensitivity to nAChR-selective antagonists but not glutamate receptor antagonists (Figure 2C). We also recorded inhibitory synaptic responses at 0 mV, which we confirmed to be mono-synaptic by the sequential addition of TTX and 4AP (Petreanu et al., 2009), and therefore not the result of feed-forward excitation mediated by ACh release (Figure 5D,E). We confirmed that these inhibitory currents were GABAergic by their sensitivity to the GABA_A_ receptor antagonist gabazine (Figure 5D,E). Notably, synaptic responses were present in only a small number of the post-synaptic neurons that we randomly recorded from near VCINs labeled with ChR2 (Figure 5F). In addition, the majority of responsive neurons showed either a cholinergic EPSC or GABAergic IPSC, and rarely both. This is consistent with a previous functional analysis of cholinergic Globus Pallidus externus connectivity to cortex (Saunders et al., 2015b), and indicates that VCINs are highly selective with the specific neuron populations they target.

**Figure 5.**
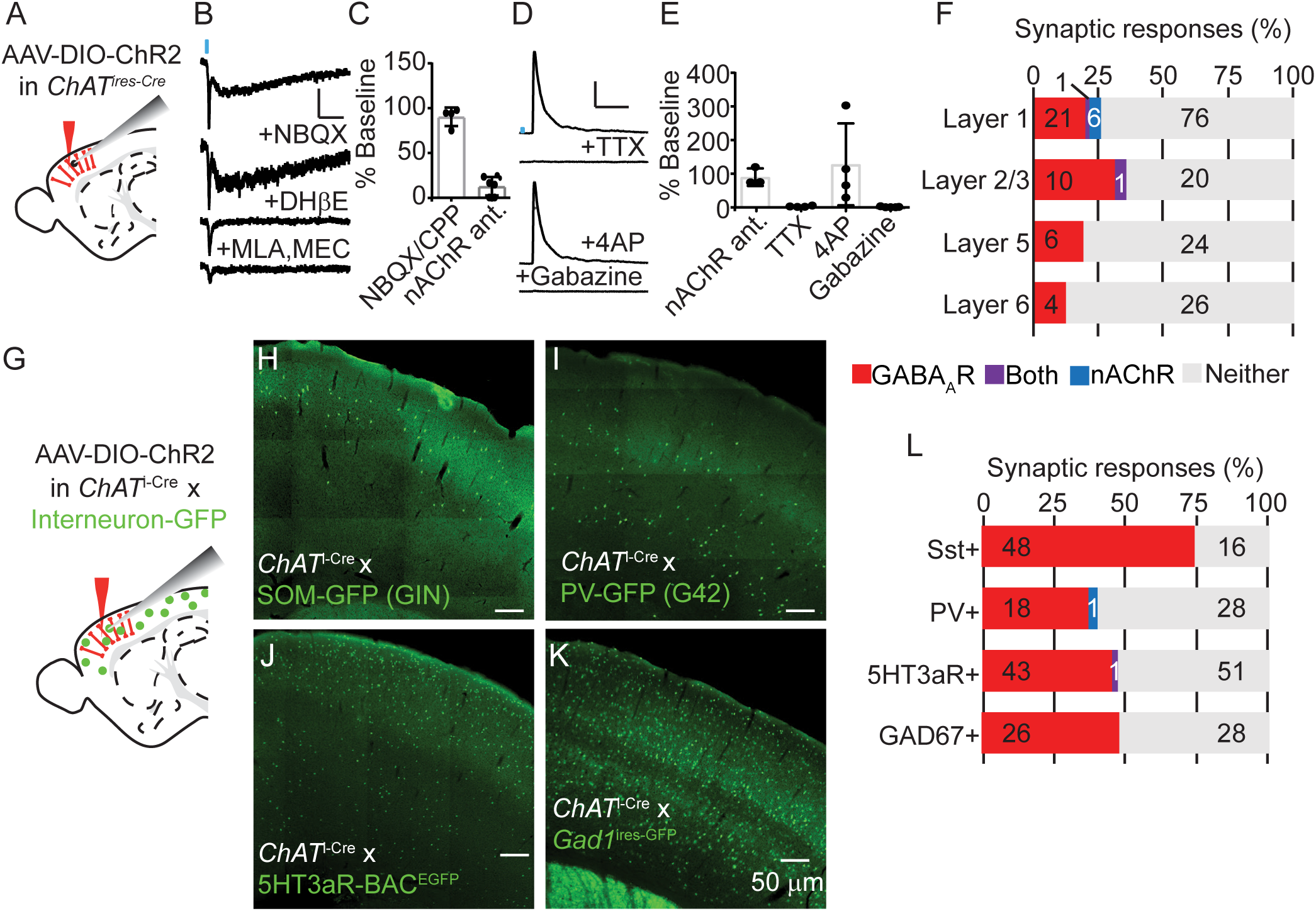
VCINs release GABA and ACh and target GABA release onto inhibitory interneurons, especially Sst+ interneurons. A) Experimental design: AAV(8)-DIO-ChR2-mCherry was injected into the frontal cortex or visual cortex of ChATires-Cre mice. Following 3-4 weeks, whole-cell voltage clamp recordings of unlabeled neurons were made from acute sagittal slices. The proportion of responses from frontal cortex and visual cortex were similar and pooled in this analysis. B) Example traces of a biphasic nAChR-mediated synaptic current recorded at -70 mV, which is insensitive to AMPA receptor antagonist NBQX. ɑ4-selective antagonist DhβE blocks the slow component, and ɑ7-selective antagonist MLA and pan-nicotinic antagonist MEC block the fast component. Scale bars represent 10 pA and 50 ms. C) Summary quantification of sensitivity to glutamatergic antagonists NBQX/CPP and nAChR antagonists DhβE, MLA, and MEC. Error bars show mean ± s.e.m. D) Example trace of a GABAA receptor-mediated synaptic currents recorded at 0 mV. Addition of TTX blocks this current, but it is rescued by subsequent addition of 4AP, indicating it is monosynaptic. Block by gabazine confirms it is mediated by GABAA receptors. Scale bars are 10 pA by 50 ms. E) Quantification of the effects of nAChR antagonists, TTX, 4AP, and gabazine on inhibitory currents elicited by stimulating VCINs. Error bars show mean ± s.e.m. F) The proportion of cells showing synaptic responses following optogenetic stimulation of VCINs. GABAAR- and nAChR-mediated responses were differentiated by clamping cells to 0 mV and -70 mV, respectively, and by response kinetics and sensitivity to channel-selective antagonists. The numbers indicate how many cells are in each category. G) Experimental design: AAV(8)-DIO-ChR2-mCherry was injected into the frontal cortex of ChATires-Cre mice crossed to different mouse lines that express GFP in specific interneuron subpopulations. (H-K) Example images showing GFP expression in 4 different mouse lines expressing GFP in different interneuron subtypes. (C) The proportion of cells of each interneuron subtype that had synaptic responses to optogenetic stimulation of VCINs. The number of cells per category are indicated.

To determine which molecularly-defined neuron subtypes receive VCIN input, we systematically surveyed the major neuronal subclasses for post-synaptic responses. To test VCIN connectivity with specific interneuron subtypes, we repeated the above ChR2-assisted connectivity survey in *ChAT^ires-Cre^* mice crossed with BAC transgenic mouse lines that express GFP in different interneuron populations, including Sst (Oliva et al., 2000), PV (Chattopadhyaya et al., 2004), and 5HT3aR (Lee et al., 2010a) expressing interneurons (Figure 5G-K). Recording from these labeled interneurons greatly increased the rate of observing GABAergic responses, especially onto Sst interneurons, and to a lesser extent PV and 5HT3aR interneurons, while ACh-mediated responses were rare (Figure 5L). Combined, these interneuron populations represent all cortical interneuron populations (Rudy et al., 2011). However, these BAC transgenic lines incompletely label their respective interneuron subpopulations, so we also recorded responses from all GABAergic interneurons labeled in GAD1^ires-GFP^ knock-in mice, and again observed only synaptic GABAergic responses (Figure 5L). For each of these interneuron populations, we confirmed that the observed GABAergic synaptic responses were monosynaptic (Figure S7). In contrast, when we specifically targeted excitatory pyramidal neurons, we observed very few GABAergic responses and no cholinergic responses (Figure S8). This is consistent with the known connectivity of VIP interneurons in general (Karnani et al., 2016a; Pfeffer et al., 2013), indicating that VCINs also mediate disinhibition of cortex.

This initial survey of VCIN connectivity shows robust and selective GABA transmission. In contrast, ACh transmission is sparse, even though 5HT3aR interneurons that we show to receive GABAergic input are known to express acetylcholine receptors (Lee et al., 2010a). This indicates that ACh transmission is highly specific, and that we likely targeted the wrong post-synaptic populations. We therefore tested the connectivity to additional post-synaptic candidates that are likely to receive cholinergic input from VCINs. A previous study has identified a subpopulation of deep layer Sst^+^ non-Martinotti interneurons that are sensitive to muscarinic receptor antagonists and increase their cortical activity in response to sensory stimulation (Muñoz et al., 2017). However, the BAC transgenic line we used to label Sst interneurons labels only a subset of superficial Sst interneurons (Figure 5H). We therefore recorded from deep-layer Sst+ cells by injecting *ChAT^ires-Cre^* x *SOM^ires-Flp^* mice with AAV(8)-DIO-ChR2-mCherry and AAV(8)-fDIO-EYFP (Figure S9A). However, we identified only GABAergic inhibitory responses in a small subset of cells (Figure S9B). Another potential target of ACh release is nAChRs on excitatory pre-synaptic terminals, as reporter by Von Engelhardt and colleagues (2007). To test this, we recorded spontaneous EPSCs (sEPSCs) from cortical pyramidal neurons while optogenetically stimulating nearby VCINs. In contrast to this previous study, we did not observe any increase in sEPSC frequency or amplitude during stimulation of VCINS (Figure S10), despite activating a much larger number of cells. This discrepancy may be explained by the use in the earlier study of a BAC transgenic line to identify the VCINs, which in a similar but different transgenic mouse drives artificially enhanced cholinergic signaling by overexpressing VAChT (Kolisnyk et al., 2013).

Another likely target of ACh release from VCINs is cortical blood vessels. ACh causes vasodilation and both VIP interneurons and cholinergic fibers abut cortical arteries (Chédotal et al., 1994; Consonni et al., 2009). Stimulation of VIP interneurons has also been shown to dilate blood vessels in acute slices (Kocharyan et al., 2008). We therefore hypothesized that ACh release from VCINs might coordinate cortical disinhibition with the neurovascular coupling necessary to supply oxygen and nutrients. Using *in vivo* imaging of pial arteries (Supplemental Fig. 11), we tested whether ACh release from VIP interneurons mediates vasodilation. We observed no difference in the vasodilation of pial arteries of mice whose VIP interneurons were genetically impaired from releasing ACh (VIP^ires-Cre^ x VAChT^fl/fl^) (Martins-Silva et al., 2011), either in response to whisker stimulation (Figure S11C) or direct optogenetic stimulation of VIP interneurons (Figure S11D).

ACh-release from VCINs may also target other VCINs. In the electrophysiological analyses described above, we stimulate the greatest possible number of local VCINs, making it impossible to record from and detect potential post-synaptic responses in the VCINs themselves. One study examining the cooperative firing of similar interneuron subtypes reported that VIP^+^ interneurons can increase each other’s action potential firing partially through activation of nAChRs (Karnani et al., 2016b). To specifically test that VCINs release ACh onto each other, we devised a strategy to robustly express ChR2 in only a subset of VCINs while labeling most with a fluorophore. We injected a diluted AAV(8)-DIO-FlpO virus, such that only a subset of VCINs were infected and activated FlpO expression. After two weeks, we injected a mixture of AAV(8)-DIO-mCherry and AAV(8)-fDIO-ChR2-EYFP. This resulted in a mixture of labeled VCINs, some of which expressed both ChR2-EYFP and mCherry, and some of which only expressed mCherry (Figure 6A). Whole-cell voltage clamp recordings from mCherry^+^/ChR2^-^ neurons while stimulating with blue-light revealed both GABA-and ACh-mediated currents, with more GABA responses than ACh responses (Figure 6B). The GABA-mediated responses were confirmed to be monosynaptic using consecutive application of TTX and 4AP (Figure 6C,D). The ACh-mediated currents were blocked by a cocktail of nAChR antagonists, and had both fast and slow kinetics, suggesting activation of both synaptic and extra-synaptic receptors (Figure 6E,F). These results confirms that VCINs transmit ACh onto interneurons of the same molecular subtype and demonstrate the remarkable specificity with which VCINs can target post-synaptic cells with different neurotransmitters.

**Figure 6.**
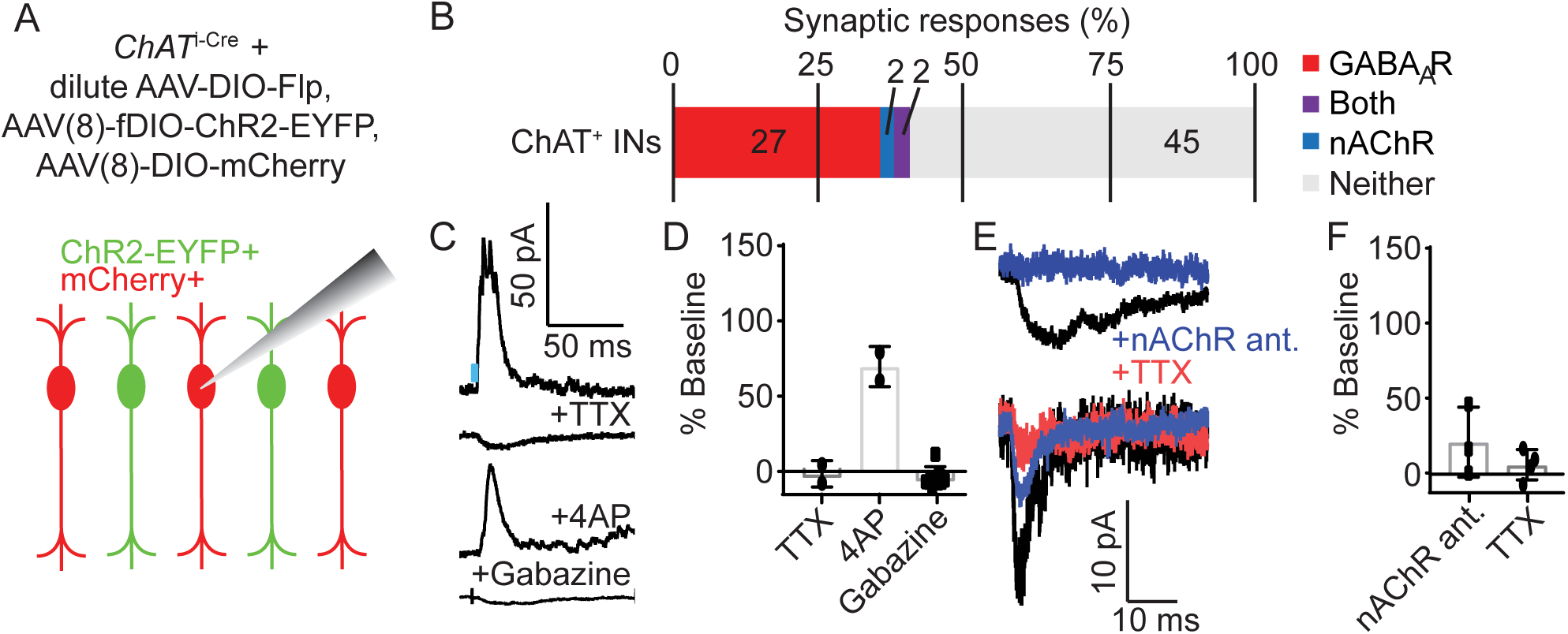
VCINs synapse onto other VCINs, releasing both ACh and GABA. A) Experimental design: To achieve mosaic expression of ChR2 in a subset of VCINs, we injected a dilute AAV(8)-DIO-FlpO virus so that a subset of VCINs would express Flp. We then injected with high-titer AAV(8)-fDIO-ChR2-EYFP and AAV(8)-DIO-mCherry. We targeted mCherry+, EYFP- cells for recording that neighbored EYFP+ neurons for whole-cell voltage clamp recording. (B) Proportion of VCINs that showed synaptic responses following stimulation of neighboring ChR2-expressin VCINs. GABAAR-mediated and nAChR-mediated responses were differentiated based on reversal potential (−70 mV and 0 mV, respectively), kinetics, and sensitivity to receptor-selective antagonists. Number of neurons in each category are indicated. (C) Example traces showing putative GABAA receptor-mediated synaptic response at baseline and following application of TTX, 4AP, and gabazine. D) Quantification of block by TTX, rescue by 4AP, and block by gabazine of putative GABAAR-mediated synaptic responses. Error bars show mean ± s.e.m. E) Example traces of two different neurons showing nAChR-mediated responses at baseline and following application of nAChR antagonists and TTX. These two cells have different response kinetics, potentially indicative of extra-synaptic (top) and synaptic (bottom) nAChRs. Scale bar: F) Quantification of putative nAChR-mediated synaptic response sensitivity to nAChR antagonists (MEC, MLA, and DHβE) and TTX.

In summary, this synaptic connectivity analysis shows that VCINs robustly release GABA onto other inhibitory interneuron subtypes, especially Sst^+^ interneurons, consistent with connectivity observed in the wider VIP^+^ interneuron population. In contrast, ACh release is sparse and selective, with the most responses observed in layer 1 interneurons (Figure 5F) and other VCINs (Figure 6). This suggests a model in which ACh and GABA co-transmission have cooperative effects, with GABA inhibiting other interneuron subtypes, and ACh reinforcing excitation of specific sub-networks of disinhibitory interneurons.

## Discussion

We describe a population of cortical VIP/ChAT interneurons that co-transmit GABA and ACh but onto different post-synaptic targets. Though underappreciated, that some VIP interneurons express ChAT has been noted for decades (Eckenstein and Baughman, 1984), can be readily identified in recently published single-cell RNA sequencing surveys (Saunders et al., 2018; Tasic et al., 2016; Zeisel et al., 2015), and is confirmed here by both immunohistochemistry and FISH. Contrary to previous reports (Bayraktar et al., 1997; von Engelhardt et al., 2007), our data establish that all VCINs are competent to release GABA. The pattern of GABA connectivity we find, with VCIN GABAergic synapses onto a diversity of interneurons, especially superficial Sst^+^ interneurons, is consistent with previous connectivity analysis of VIP interneurons (Karnani et al., 2016a; Pfeffer et al., 2013). This does not preclude further functional heterogeneity within VIP interneurons, as demonstrated by evidence that VIPs can directly inhibit excitatory pyramidal neurons (Garcia-Junco-Clemente et al., 2017; Zhou et al., 2017).

Compared to GABA release from VCINs, ACh release is even more specific and targeted. Given the post-synaptic ACh-mediated currents we observed combined with the evidence from our molecular analyses, it is clear that these neurons are indeed competent to release ACh. The subtypes of neurons that we observed to receive cholinergic input from VCINs, layer 1 interneurons and the VCINs themselves, are consistent with ACh release providing a net excitatory effect to complement disinhibition mediated by GABA. While we endeavored to be as comprehensive as possible in surveying potential post-synaptic targets, the full diversity of cortical cellular subtypes is only beginning to be understood, and the tools to target those various subtypes selectively are still being developed. It is therefore possible that future research will more definitively identify a neuronal subtype that is strongly innervated by ACh from VCINs. It is also possible that ACh release is regulated such that it only occurs in certain contexts or developmental epochs. Regulation of neurotransmitter release has been observed in other systems, such as the retina where ACh and GABA are differentially released by starbust amacrine cells depending on the direction of the light stimulus (Lee et al., 2010b; Sethuramanujam et al., 2016), and in several examples of neurons that appear to switch their predominant neurotransmitter during developing or after bouts of neuronal activity (Spitzer, 2017). In any case, ACh transmission from VCINs is finely controlled, such that the majority of post-synaptic neurons do not receive cholinergic input.

This level of control is highlighted by the specialization of VCIN pre-synaptic terminals, where only a subset is competent to release ACh. Such output-specific targeting of neurotransmitter release is a largely unexplored aspect of synaptic transmission in the brain. A similar level of regulation can be observed in dopaminergic neurons which spatially segregate co-release of glutamate and dopamine in different brain regions (Mingote et al., 2015; Stuber et al., 2010), and whose individual axons segregate terminals that release dopamine or glutamate (Zhang et al., 2015). Similar differentiation of neurotransmitter release has been reported elsewhere in the cholinergic system, specifically in Globus Pallidus externus projections to the cortex (Saunders et al., 2015b) and in hippocampus-projecting septal cholinergic neurons that release ACh and GABA from different synaptic vesicles (Takács et al., 2018). The possibility for separable release of multiple neurotransmitters adds another level of complexity to our understanding of how neurons communicate.

The pattern of GABA and ACh connectivity we observed suggests a coherent model of the net effect that these neurons have on cortical circuits: ACh-mediated excitation of specific layer 1 interneurons and other VCINs, combined with the GABA-mediated disinhibition provides a powerful activating signal to local cortical areas. In different behavioral paradigms, activation of VIP^+^ and layer 1 interneurons has been shown to increase the gain of sensory responses in pyramidal neurons (Fu et al., 2014; Pi et al., 2013) or signal a cue for fear conditioning (Letzkus et al., 2011). In each of these cases, these disinhibitory neurons are activated by ascending cholinergic inputs from basal forebrain. We propose that ACh release from VCINs reinforces and amplifies the cortical activation achieved by the broader VIP^+^ interneuron population, itself activated by ascending cholinergic projections, in order to enhance the response to sensory cues. However, we also find VCIN-mediated IPSCs in these same ACh-sensing populations, suggesting additional complexities. One possibility is that there is spatial precision in the targeting of ACh and GABA . For example, VCINs might target other VCINs and layer 1 interneurons within the same cortical column with ACh but those in neighboring columns with GABA.

VCINs are notable not only because they co-transmit two small molecule neurotransmitters with opposite ionotropic valences, but also for their local release of ACh. Typically, neuromodulators like ACh are released by long-range projections originating from well-defined basal nuclei. Because basal forebrain cholinergic neurons send long-range and highly ramifying projections to large areas of cortex, modulation by ACh is thought to influence entire cortical regions. Highly specific ACh release by cortical interneurons suggests that neuromodulation by ACh can be targeted in small areas of cortex onto specific post-synaptic cell types. This is consistent with a recent reevaluation of cortical ACh signaling as not only a diffuse, tonic signal that operates on relatively slow time scales, but also as a phasic, point-to-point signal that operates on the time scale of seconds and milliseconds (Sarter et al., 2014).

The existence of VCINs also requires a reevaluation of studies that globally manipulate cholinergic signaling in cortex. While many studies specifically targeted cortically-projecting basal forebrain neurons, several have used genetic crosses that affect all cholinergic neurons in the brain (Chen et al., 2015; Dasgupta et al., 2018; Kuchibhotla et al., 2017; Sparks et al., 2018), and therefore include confounding effects from local VCINs. In addition, studies that use ChAT-BAC-ChR2 mice to activate cholinergic neurons not only run the risk of confounding effects from overexpressed VAChT (Kolisnyk et al., 2013), but also from incidental manipulation of cortical VIP interneurons, which are known to have profound effects on cortical function even purely through GABA release. Going forward, studies of cholinergic signaling in cortex must differentiate between contributions from basal forebrain projections and those from local cholinergic interneurons.

## Methods

### Mice

We used the following mouse lines in this study:

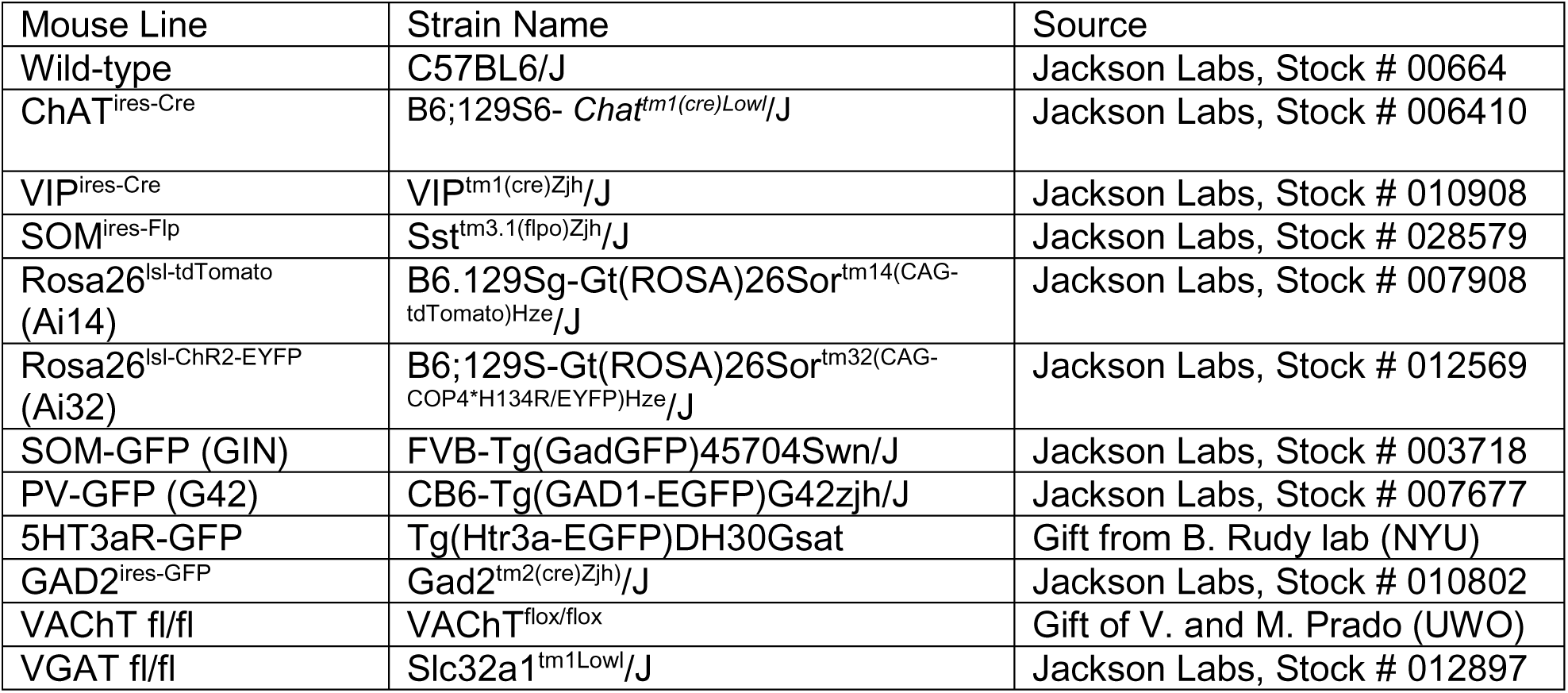

| Mouse Line | Strain Name | Source |
| --- | --- | --- |
| Wild-type | C57BL6/J | Jackson Labs, Stock # 00664 |
| ChAT<sup>ires-Cre</sup> | B6;129S6- *Chat*<sup>tm1(cre)Lowl/J</sup> | Jackson Labs, Stock # 006410 |
| VIP<sup>ires-Cre</sup> | VIP<sup>tm1(cre)Zjh/J</sup> | Jackson Labs, Stock # 010908 |
| SOM<sup>ires-Flp</sup> | Sst<sup>tm3.1(flpo)Zjh/J</sup> | Jackson Labs, Stock # 028579 |
| Rosa26<sup>Isl-tdTomato</sup><br>(Ai14) | B6.129Sg-Gt(ROSA)26Sor<sup>tm14(CAG-tdTomato)Hze/J</sup> | Jackson Labs, Stock # 007908 |
| Rosa26<sup>Isl-ChR2-EYFP</sup><br>(Ai32) | B6;129S-Gt(ROSA)26Sor<sup>tm32(CAG-COP4*H134R/EYFP)Hze/J</sup> | Jackson Labs, Stock # 012569 |
| SOM-GFP (GIN) | FVB-Tg(GadGFP)45704Swn/J | Jackson Labs, Stock # 003718 |
| PV-GFP (G42) | CB6-Tg(GAD1-EGFP)G42zjh/J | Jackson Labs, Stock # 007677 |
| 5HT3aR-GFP | Tg(Htr3a-EGFP)DH30Gsat | Gift from B. Rudy lab (NYU) |
| GAD2<sup>ires-GFP</sup> | Gad2<sup>tm2(cre)Zjh/J</sup> | Jackson Labs, Stock # 010802 |
| VACHT<sup>fl/fl</sup> | VACHT<sup>flox/flox</sup> | Gift of V. and M. Prado (UWO) |
| VGAT<sup>fl/fl</sup> | Slc32a1<sup>tm1Lowl/J</sup> | Jackson Labs, Stock # 012897 |

All mice used in this study were between 2-4 months in age. For experiments using only ChAT^ires-Cre^ mice, homozygous mice were maintained. For all crosses of two mouse lines, homozygous breeders were used to produce heterozygous off-spring for experiments, with the exception of experiments requiring conditional deletion of VGAT or VAChT, in which case homozygous VGAT^fl/fl^ or VAChT^fl/fl^ conditional knock-out mice were produced that were either homozygous or heterozygous for ChAT^ires-Cre^ or VIP^ires-Cre^, respectively. All experiments were performed according to animal care and use protocols approved by the Harvard Standing Committee on Animal Care in compliance with guidelines set for in the NIH *Guide for the Care and Use of Laboratory Animals*.

### Virus Injections

For intracranial injection of virus, the surgery work area was maintained in asceptic conditions. Mice were anesthetized with 2-3 % isoflurane and given 5 mg/kg ketoprofen as prophylactic analgesic, and placed on a heating pad in a stereotaxic frame (David Kopf Instruments) with continuous delivery and monitoring of appropriate isoflurane anesthesia. The skin above the skull was carefully cleared of hair with scissors and depilatory cream (Nair) and sterilized with alternating scrubs with alcohol pads and betadine pads. A midline incision was made in the skin and the skull exposed. Small holes were drilled into the skull at stereotactic coordinates ± 1.8 mm ML, + 1.8 mm and + 0.5 mm AP from bregma. In a subset of experiments, we targeted visual cortex by injecting at stereotactic coordinates ± 2.5 mm and 0 mm from lambda. 200-500 μl of the appropriate virus was injected through a pulled glass pipette at a rate of 100 nl/min with a UMP3 microsyringe pump (World Precision Instruments) at a depth of -0.6 mm below the pia, or -0.25 mm below pia for visual cortex. Following injection, the pipette was allowed to sit for 10 minutes to prevent leak of the virus from the injection site, and then the glass pipette slowly removed over the course of 1-2 mintues. Following surgery, mice were monitored in their home cage for 4 days following surgery, and received daily analgesia for 2 days following surgery. Mice were sacrificed for experiments at least 3 weeks following injection to allow for robust viral expression. When we injected multiple viruses, they were mixed in equal proportions. The following viruses were used in this study:

| Virus | Titer | Figures |
| --- | --- | --- |
| AAV(8)-EF1α-hChR2(H134R)-mCherry-WPRE-pA | $\sim 2 \times 10^{13}$ gc/ml | Fig. 5,6; Sup Fig. 7,8,10,11,12 |
| AAV(DJ)-EF1α-fDIO-EYFP | $\sim 2 \times 10^{12}$ gc/ml | Sup Fig. 10 |
| AAV(1)- EF1α-fDIO-ChR2-EYFP | $\sim 5 \times 10^{13}$ gc/ml | Fig. 6 |
| AAV(8)- EF1α-DIO-FlpO | $\sim 7 \times 10^{11}$ gc/ml | Fig. 6 |
| AAV(8)- EF1α-DIO-mCherry | $\sim 6 \times 10^{12}$ gc/ml | Fig. 6 |
| AAV(8)-CMV-DIO-Synaptophysin-EYFP | $\sim 6 \times 10^{12}$ gc/ml | Fig. 2,3; Sup Fig 4,5 |
| AAV(9)-CAG-DIO-Synaptophysin-mCherry | $\sim 2 \times 10^{13}$ gc/ml | Fig. 4; Sup Fig 6 |

### Electrophysiology

300 µm acute coronal brain slices were prepared from mice deeply anesthetized with isoflurane inhalation and perfused with ice-cold cutting solution containing (in mM): 25 NaHCO_3_, 25 Glucose, 1.25 NaH_2_PO_4_, 7 MgCl_2_, 2.5 KCl, 0.5 CaCl2, 11.6 ascorbic acid, 3.1 pyruvic acid, 110 Choline chloride. Following dissection, brains were blocked by cutting along the mid-sagittal axis, and brains glued to the platform along the mid-sagittal surface before slicing on a Leica VT1000s vibratome, while maintaining submersion in cold choline cut solution. Following cutting, slices recovered for 30-45 minutes in 34° C artificial cerebral spinal fluid (aCSF) containing (in mM): 125 NaCl, 2.5 KCl, 1.25 NaH_2_PO_4_, 25 NaHCO_3_, 11 glucose, 2 CaCl_2_, 1 MgCl_2_. Subsequently all recording took place in continuous perfusion (2-3 ml/min) of room temperature aCSF. Both the choline cut solution and aCSF were continuously equilibrated by bubbling with 95% 0_2_/5% CO_2_.

After recovery, slices were transferred to a recording chamber mounted on an upright microscope (Olympus BX51WI). Cells were imaged using infrared-differential interference contrast with a 40x water-immersion Olympus objective. To confirm ChR2 expression and GFP-labeled interneurons, we used epifluorescence with an X-Cite 120Q (Excelitas) as a light source. Whole cell voltage-clamp and current clamp recordings were obtained by forming intracellular seals with target neurons with patch pipettess pulled from borosilicate glass (BF150-86-7.5, Sutter). Pipettes (2-4 MOhm pipette resistance) were pulled with a P-97 flaming micropipette puller (Sutter). Pipettes were filled with either a Cs^+^-based internal recording solution containing (in mM): 135 CsMeSO_3_ 10 HEPES, 1 EGTA, 4 Mg-ATP, 0.3 Na-GTP, 8 Na_2_-Phosphocreatine, 3.3 QX-314 (Cl- salt), pH adjusted to 7.3 with CsOH and diluted to 290-295 mOsm/kg for voltage clamp recordings or a K^+^-based internal recording solution containing (in mM): 120 KMeSO_3_, 10 HEPES, 0.2 EGTA, 8 NaCL, 10 KCL, 4 Mg-ATP, 0.3 Na-GTP, pH adjusted to 7.3 with CsOH and diluted to 290-295 mOsm/kg for current clamp recordings.

To stimulate ChR2-expressing neurons, we focused a fiber-coupled 200 mW 473 nm laser (Opto-engine) onto the back aperture of 40x Olympus objected in the imaging path. Laser intensity was adjusted using a neutral density filter such that ~ 9 mW/mm^2^ of total light reached the slice. ChR2^+^ cells were regularly patched to confirm that laser intensity was well above the threshold needed to elicit action potentials at low latency (data not shown). Cells were classified as having a synaptic response based on the average of at least 10 individual sweeps of optogenetic stimulation. If a consistent, time-locked response above the baseline noise could be observed, additional sweeps were taken to get a more accurate representation of the response size and kinetics. Putative GABA-mediated currents were isolated by voltage clamping at 0 mV, the reversal potential for excitatory currents, or by identified in current clamp as hyperpolarizing potentials, and confirmed with 10 µM Gabazine (SR-95531, Tocris). Putative ACh-mediated currents were isolated by voltage clamping at -70 mV, the reversal potential for inhibitory currents, or in current clamp as depolarizing potentials, and confirmed with 10 µM Methyllycaconitine citrate (MLA, Tocris), which is selective for alpha7-containing nicotinic receptors, 10 µM Dihydro-beta-erythroidine hydrobromide (DHBE, Tocris), which is selective for alpha4-containing nicotinic receptors, and 10 µM Mecamylamine hydrochloride (MEC, Tocirs), which is a non-selective nicotinic receptor antagonist. To confirm monosynaptic release of GABA, we consecutively added 1 µM TTX (AbCam) followed by 100 µM 4-aminopyridine (Tocris). To rule out contributions from other low latency excitatory receptors, we also added the glutamate receptor antagonists NBQX and CPP (both 10 µM, Tocris). For current-clamp recordings with putative muscarinic receptor-mediated currents, we washed on 10 µM Scopolamine hydrobromide (Tocris).

Voltage clamp and current clamp recordings were amplified and filtered at 3 kHz using a Multiclamp 200B (Axon Instruments) and digitized at 10 kHz with a National Instruments acquisition boards. 3 kHz. Data was saved with a custom version of ScanImage written in Matlab (Mathworks; https://github.com/bernardosabatinilab/SabalabSoftware_Nov2009). Additional off-line analysis was performed using Igor Pro (Wavemetrics). Response amplitudes were determined by averaging 5-10 traces, taking a 990 ms baseline prior to stimulation, and subtracting that from the peak amplitude within 5-20 ms after stimulation.

### Fluorescent In situ hybridization

Whole brains dissected from deeply anesthetized wild-type C57/BL6 mice were fresh frozen in Tissue-tek OCT media on dry ice and stored at -80 °C before being sliced into 20 µm slices on a CM 1950 Cryostat (Leica), mounted on SuperFrost Plus 25 x 75 mm slides (VWR), and stored at -80 °C prior to labeling. Fluorescent in situ hybridization labeling was perfomed according to the RNAscope Fluorescent Multiplex Assay protocol (ACDBio). We used the following probes from ACDBio:

| Probe name | Gene | Protein | Catalog # |
| --- | --- | --- | --- |
| Mm-Chat-C2 | *ChAT* | ChAT | 408731-C2 |
| Mm-Slc32a1 | *Slc32a1* | VGAT | 319191 |
| Mm-Gad1-C3 | *Gad1* | GAD67 | 400951-C3 |
| Mm-Gad2-C3 | *Gad2* | GAD65 | 439371-C3 |
| Mm-Slc5a7-C3 | *Slc5a7* | ChT | 439941-C3 |
| Mm-Slc18a3-C3 | *Slc18a3* | VACHT | 448771-C3 |
| Mm-Vip | *Vip* | VIP | 4159341 |
| Cre-01-C3 | *Cre* | Cre | 474001-C3 |

### Immunohistochemistry

Tissue for immunostaining was obtained from deeply anesthetized mice that were perfused transcardially with room temperature phosphate-buffered saline (PBS) followed by 4% paraformaldehyde (PFA) in PBS. The brain was then dissected out of the skull, post-fixed overnight at 4 °C in 4% PFA, rinsed and stored in PBS. Brains were sliced into either 50 µm (for most figures) or 25 slices µm (for Fig. 6) on a Leico VT1000s vibratome and stored in 24-well plates.

For staining, slices were first incubated in blocking buffer (10% Normal Goat Serum, 0.25% Triton-X in PBS, except 10% Normal Horse Serum for ChAT immunostaining) for 1 hour at room temperature on a rotary shaker, then placed in primary antibody solution (1:500 for each primary antibody diluted into carrier solution (10% Normal Goat Serum, 0.2% Triton-X in PBS) and left to shake overnight at 4 °C. Slices were then washed 5-6 x in PBS, and placed into secondary antibody solution (1:500 in carrier solution) for 2 hours at room temperature. Slices were again washed, placed on glass slides, and mounted in Prolong Gold antifade mounting media with DAPI (Invitrogen).

### Imaging and analysis

Immunostained and FISH sample slides were initially imaged on a VS120 slide scanner at 10x. Regions of interest were then imaged on either a FV1200 confocal microscope (Olympus) or a TCS SP8 confocal microscope (Leica) for colocalization analysis.

Immunostained samples were manually scored by a researcher to count co-labeled cells using the Cell Counter plugin in Fiji (https://fiji.sc/). FISH samples were analyzed with an automated analysis pipeline custom written using Fiji and Matlab. A cellular mask was created by combining the 3 FISH channels and using the Renyi entropy thresholding algorithm to binarize the image. Each individual cell was identified, and the percent coverage of each FISH channel was calculated for each cell. A threshold to classify each cell as positive or negative for each FISH channel was then determined by selecting a threshold for percent coverage above ten manually-drawn background areas. From this analysis, we determined the proportion of cells that are positive for each of the different FISH probes.

Immunostained samples from Figure 6 were imaged using the TCS SP8 confocal microscope (Leica) such that each acquisition utilized the full dynamic imaging range. For analysis, putative individual pre-synaptic terminals were identified by thresholding the raw image stacks of the Synaptophysin-mCherry signal, then filtering putative terminals for size and enforcing that they must be present across multiple images. Mean fluorescence intensity for VGAT and VAChT antibody staining was calculated for each putative terminals. Individual terminals were classified as VGAT or VAChT positive by automatically determining a threshold for VGAT/VAChT positive pixels using the Otsu method (which determines the intensity threshold that minimizes intraclass variance and maximizes interclass variance), and requiring that the terminal is positive or negative if the mean intensity is greater than or equal to the Otsu threshold.

### Array Tomography

For array tomography, brains from mice injected with AAV(8)-CMV-DIO-Synaptophysin-YFP were perfused, dissected, and fixed as for immunohistochemistry. 300 µm thick slices were then cut with a Lieca VT1000s vibratome. Areas of high Synaptophysin-YFP expression were noted using an epifluorescence microscope, and approximately 1 x 1 mm squares of tissue were cut out under a dissecting scope with Microfeather disposable ophthalmic scalpals. These small tissue squares were then dehydrated with serial alcohol diluations and infiltrated with LR White acrylic resin (Sigma Aldrich L9774), and placed in a gel-cap filled with LR White to polymerize overnight at 50 °C. Blocks of tissue were sliced on an ultramicrotome (Leica EM UC7) into ribbons 70 nm sections.

Antibody staining of these sections were performed as previously described (Saunders et al., 2015b). Breifly, antibodies were stained across multiple staining sessions, with up to three antibodies stained per session, and a fourth channel left for DAPI. Typically, Session 1 stained against YFP (chicken ɑ-GFP, GTX13970, GeneTex), Gephyrin (mouse ɑ-Gephyrin, 612632, Biosciences Pharmingen), and Synapsin-1 (rabbit ɑ-Synapsin-1, 5297S, Cell Signaling Tech), Session 2 for PSD-95 (rabbit ɑ-PSD95, 3450 Cell Signaling Tech.), Session 3 for VGAT (rabbit ɑ-VGAT, 131 011 Synaptic Systems), Session 4 for VAChT (mouse ɑ-VAChT, 139 103 Synaptic Systems) and VGLUT1 (guinea pig ɑ-VGAT, AB5905 Millipore), and Session 5 for ChAT (goat ɑ-ChAT, AB144P Millipore). One test sample was performed where the staining order was reversed, and while staining quality did appear degraded for later samples, it was not significant enough to alter analysis. Each round of staining was imaged on a Zeiss Axio Imager upright fluorescence microscope before the tissue ribbons were stripped of antibody and re-stained for a new session of imaging. Four images were acquired with a 63x oil objective (Zeiss) and stitched into a single final image (Mosaix, Axiovision). Image stacks were processed by first aligning in Fiji with the MultiStackReg plug-in, first on the DAPI nuclear stain, with fine alignments performed using the Synapsin 1 stack. Fluorescence intensity was also normalized across all channels, such that the top and bottom 0.1% of fluorescence intensities were set to 0 and maximum intensity, respectively.

For analysis, Synaptophysin-YFP masks were created by first masking out the edges of the images that did not contain any tissue sample and the DAPI signal to exclude cell nuclei, then by empirically determining an appropriate threshold of YFP fluorescence. Putative pre-synaptic terminals were required to exist on multiple z-places of the image stack, thus creating 3D binary masks corresponding to putative pre-synaptic terminals. Global cross-correlations were made by z-scoring the fluorescence signals of each antibody stack making pairwise comparisons among all stacks, shifting the images +/- 10 pixels vertically and horizontally and calculating the 2D co-variance at every shift. The area of the images covered by VCIN-expressed Synaptophysin-YFP is ~0.1% of the total – thus the co-expression of synaptic markers within these terminals contributes minimally to the global cross-correlations reported above. To avoid amplifying any small background signals that would result if an antibody signal was low in the YFP^+^ pre-synaptic terminals, we z-scored the fluorescence intensities across the entire image stack (as for the global cross correlation analysis above) but calculated the co-variance across signal pairs only within the YFP^+^ terminals.

Colocalization analysis was carried out using the same YFP mask as described above. Synaptic antibody signals was assigned to individual pixels by fitting each antibody punctum with a Gaussian distribution, and assigning the pixel corresponding to the peak of that Gaussian as the location of that antibody. Colocalization was then calculated by dividing the number of antibody pixels that overlapped with the YFP mask by the total number of pixels in the YFP mask. Similar colocalization values were also calculated within expanding single-pixel concentric volumes around each terminal, to compare the antibody colocalization within terminals with the immediately surrounding tissue. Finally, the location of each antibody puncta was randomized 1000 times, avoiding the DAPI masks, and the colocalization within and around the YFP terminals recalculated for each round of randomization. To compare across samples, this colocalization measure was converted to a z-score by subtracting the mean of the randomized data from the actual colocalization, divided by the standard deviation of the randomized data.

### Blood vessel imaging

For surgical implantation of cranial windows, mice were anesthetized with 2-3% isoflurane, given 10 mg/kg ketoprofen as prophylactic analgesic, and 0.3 mg/kg Dexamethasone to limit tissue inflammation. Mice were placed on a heating pad in a stereotaxic frame (David Kopf Instruments) with continuous delivery and monitoring of appropriate isoflurane anesthesia. The skin above the skull was carefully cleared of hair with scissors and depilatory cream (Nair) and sterilized with alternating scrubs with alcohol pads and betadine pads. A midline incision was made in the skin and the skull exposed. A circular, ~3 mm diameter section of skull was carefully drilled from over the right barrel cortex, with frequent application of sterile saline. A cranial window, prepared by adhering a 3 mm glass coverslip to a 4 mm coverslip with optical glue, was placed over the brain, and secured in place with Kwik-cast silicone elastomer sealant (World Precision Instruments), followed by C&BMetabond (Parkell) with a custom-made titanium head post. Following surgery, mice were monitored in their home cage for 4 days following surgery, and received daily analgesia for 2 days following surgery

Alexa Fluor 633 hydrazide (5 mg•kg^-1^) was retro-orbitally injected into mice to visualize arterioles *in vivo.* Arterioles were imaged at 800 nm with a field of view size of 200 µm x 200 µm (512×512 pixels, pixel size of 0.16 µm^2^/pixel) at 30 Hz. Optical stimulation was performed using pulsed illumination (5 pulses, 20 Hz, 5 ms ON/45 ms OFF, 30 mW/mm2) using a 473 nm solid-state laser. Whisker stimulation (4 Hz, 5 s) was performed using a foam brush controlled by a servo motor under the control of Wavesurfer. Three technical trials where acquired and averaged for each field of view. 10-13 fields of view were acquired per imaging session. Three imaging sessions were collected on three separate days per mouse and arteriolar dilation responses were averaged across all three sessions for each mouse.

## Author Contributions

A.J.G. and B.L.S. conceived the study. W.W., and A.S. collected and analyzed electrophysiology data. K.R., A.Z., and K.B. collected immunohistochemistry and *in situ* data. B.C., V.N., and C.G. collected and analyzed *in vivo* imaging of cortical blood vessels. A.J.G. was involved in collection and analysis of all experiments. A.J.G. and B.L.S. wrote the manuscript with comments and feedback from the other authors.

## Acknowledgements

The authors thank V. Prado and M. Prado for generous donation of the VAChT^fl/fl^ mouse line, B. Rudy for donation of the 5HT3aR-GFP mouse line, and M. El-Rifai, N. Kingery, and T. Xie for production and analysis of array tomography data. Thanks to members of the Sabatini lab for thoughtful critique of the manuscript. This work was supported by a fellowship from Jane Coffin Childs Fund (A.J.G), and grants from the NIH (K99 NS102429 to A.J.G., R37 NS046579 to B.L.S., and P30NS072030 to the Neurobiology Imaging Facility).

## Supplemental Figures

**Figure S1.**
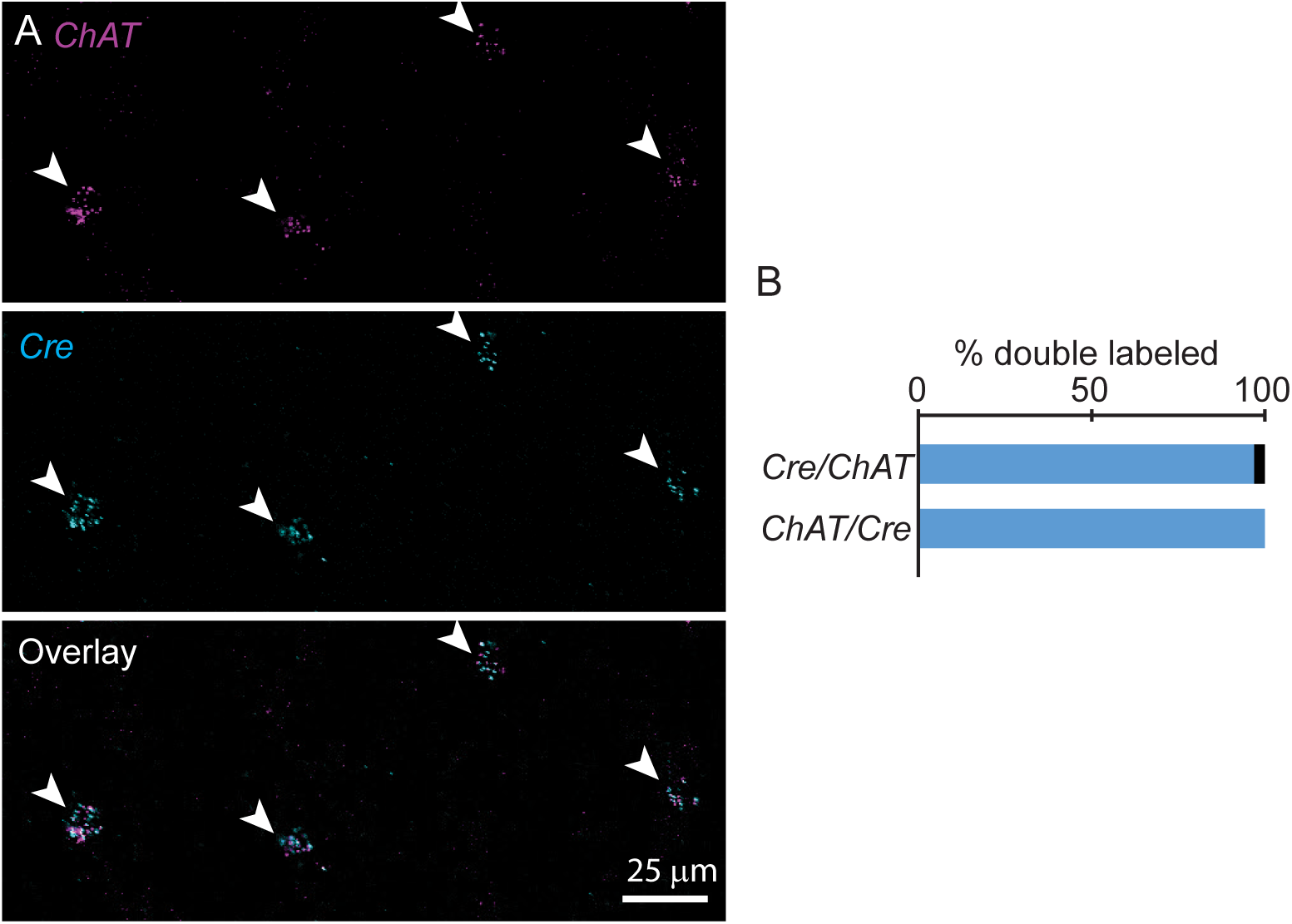
Chat^ires-Cre^ mice faithfully express cre only in ChAT+ neurons in the cortex. A) Example fluorescent in situ hybridization of ChAT (top), Cre (middle), and the overlay (bottom) from ChAT^ires-Cre^ mice. Arrow heads indicate dual ChAT+, Cre+ cells. B) Quantification of cells expressing both ChAT and Cre (n = 32 ChAT+,Cre+ of 33 ChAT+ and 32 Cre+ neurons).

**Figure S2.**
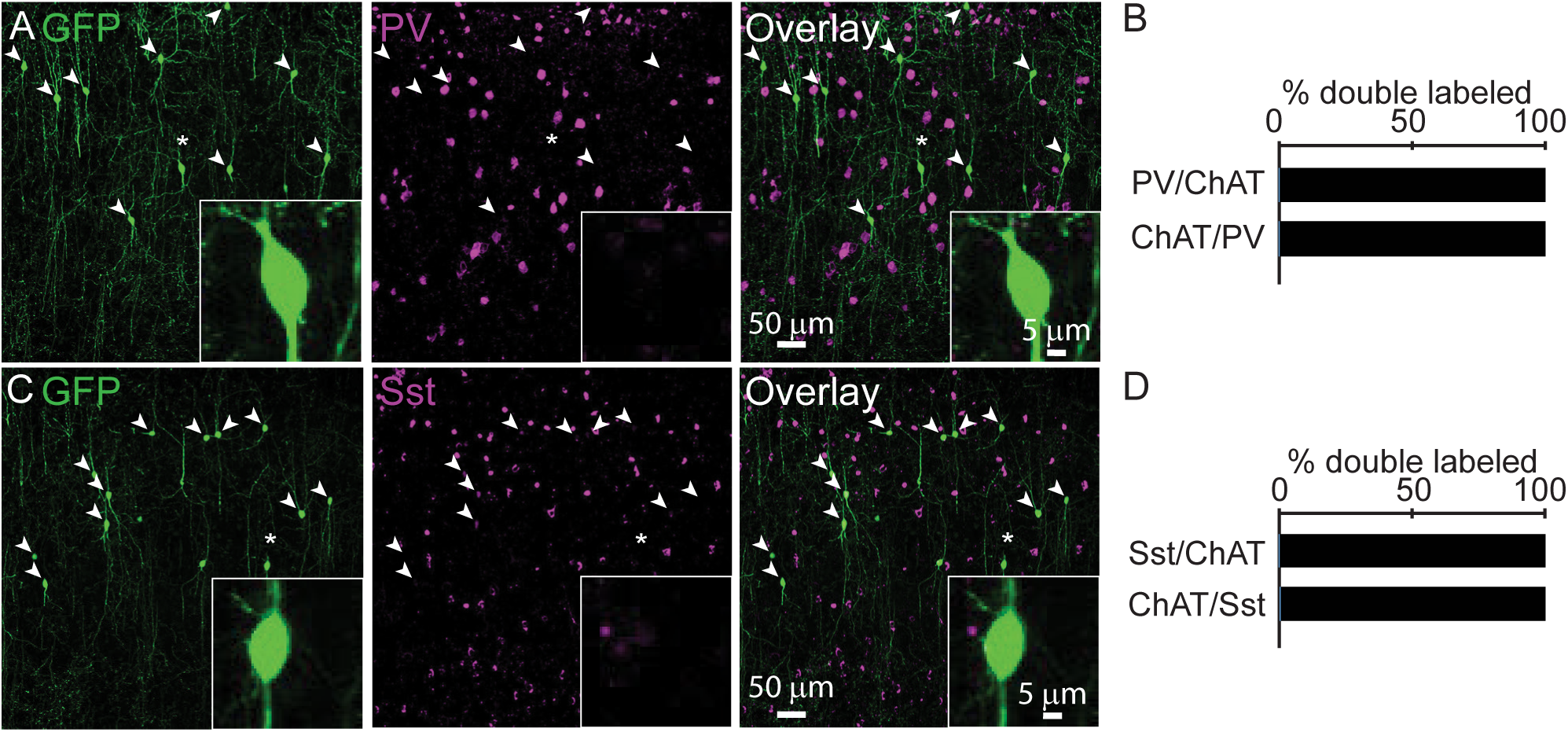
Cortical ChAT+ neurons do not express Parvalbumin or Somatostatin. A,B) Example images (left) and quantification (right) of cortical ChAT+ neurons (left panel), labeled by injecting AAV(8)-DIO-EGFP into the frontal cortex of ChATires-Cre mice. Immunostaining against parvalbumin (PV, middle panel) shows no co-labeling with cortical ChAT+ neurons (overlay, right panel; n = 1 ChAT+,PV+ of 180 CHAT+ and 576 PV+ neurons). C,D) Example images (left) and quantification (right) of GFP-labeled cortical ChAT+ neurons (left panel) and immunostaining against Somatostatin (Sst, middle panel) showing no co-labeling (overlay, right panel; n = 2 ChAT+,Sst+ of 360 ChAT+ and 1016 Sst+ neurons). Arrowheads indicate GFP-expressing ChAT+ neurons and asterisks indicate cells shown in the insets.

**Figure S3.**
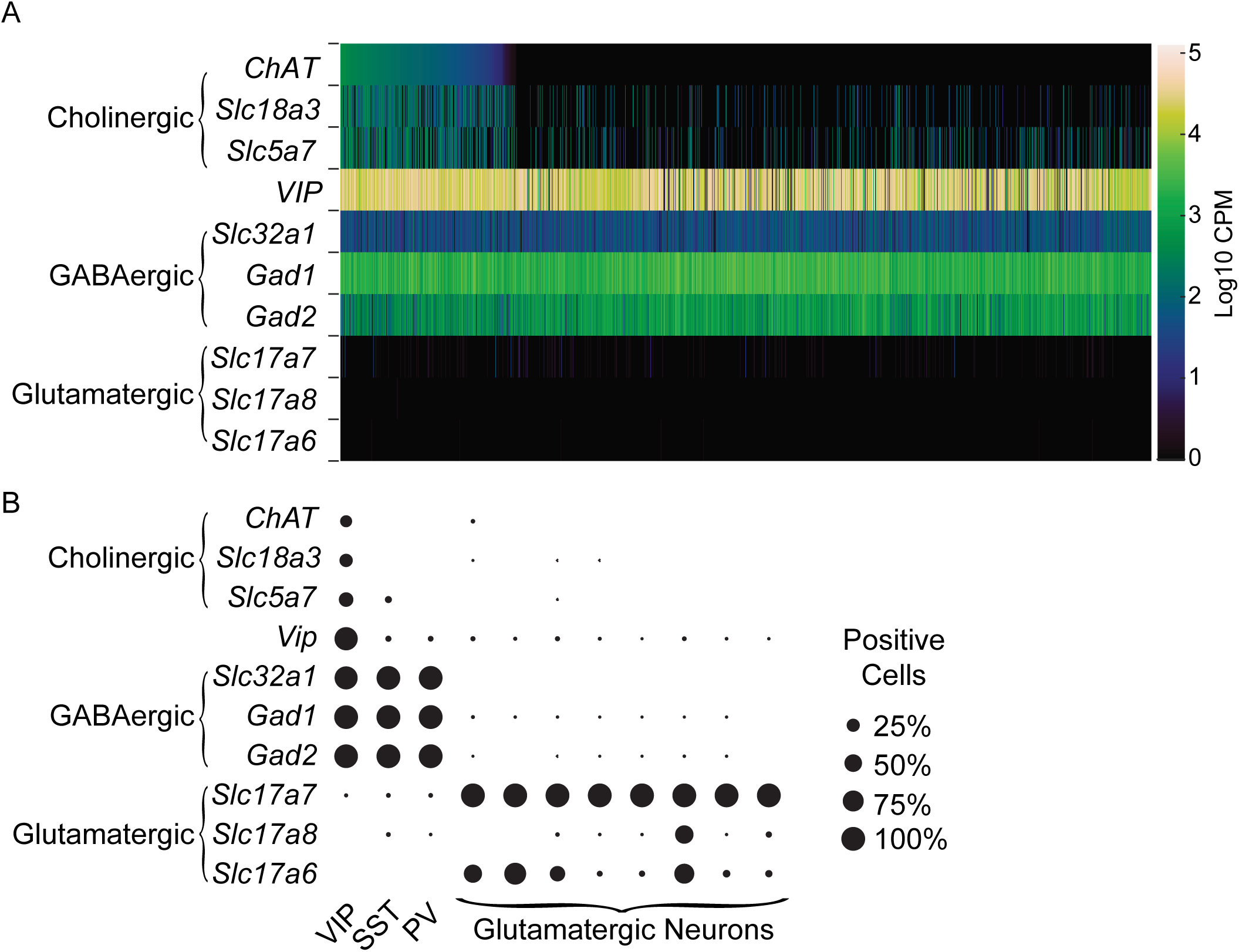
A subset of VIP interneurons express cholinergic genes by single-cell RNA sequencing. A) A heat map of the number of transcripts per cell (log counts per million, Log10 CPM) for various neurotransmitter synthesis and vesicular release machinery genes from 2,952 single-cell transcriptomes of VIP-expressing neurons from visual cortex and anterior lateral motor cortex. Data is downloaded from the Allen Brain Institute’s RNA-Seq Data Navigator, accessible at: http://celltypes.brain-map.org/rnaseq/mouse. (B) The proportion of cells in different cell sub-types that are positive for different transcripts for genes indicating different neurotransmitter phenotypes, as defined by > 1 log10 CPM of each transcript.

**Figure S4.**
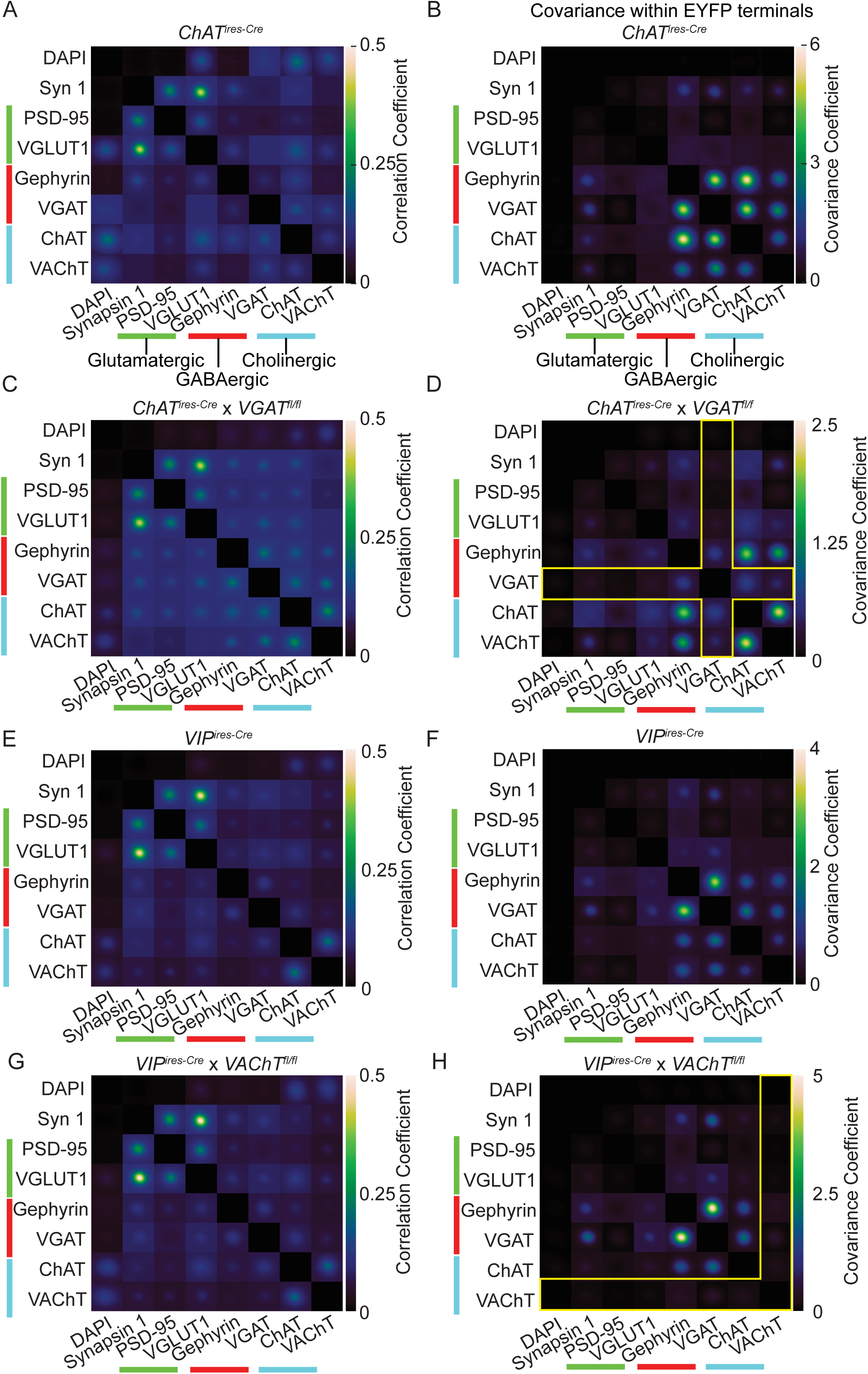
Average cross-correlations of synaptic antibody image arrays from ChAT^ires-Cre^, VIP^ires-Cre^, VGAT mosaic conditional knock-out (ChAT^ires-Cre^ x VGAT^fl/fl^) and VAChT mosaic conditional knockout mice (VIP^ires-Cre^ x VAChT^fl/fl^). A,B) Global cross-correlations (A) and co-variances within YFP masks of putative VCIN pre-synaptic terminals (B) of tissue collected from ChATires-Cre mice injected with AAV(8)-DIO-Synaptophysin-YFP. These graphs are duplicated from Figure 2D & E. C,D) To test the specificity of the VGAT antibody for VCIN terminals, we repeated the analysis from Fig. 2 in tissue from ChAT^ires-Cre^ x VGAT^fl/fl^ mice (Tong et al., 2008), and found that VGAT staining is no longer correlated with Gephyrin or the cholinergic markers. Yellow bars in (D) highlight the loss of VGAT correlations within VCIN terminals compared to (B). E,F) To confirm the specificity of VAChT staining, we selectively deleted VAChT in VIP interneurons (VIP^ires-Cre^ x VAChT^fl/fl^). We therefore first tested the global cross-correlations and YFP-masked co-variances of tissue collected from VIPires-Cre mice injected with AAV(8)-DIO-Synptophysin-YFP, and saw similar patterns of enrichment for GABAergic and Cholinergic markers as in (B). G,H) In contrast to tissue from VIP^ires-Cre^ mice, global cross-correlations and YFP-masked co-variances of tissue collected from mice with VAChT deleted from all VIP interneurons (VIP^ires-Cre^ x VAChT^fl/fl^) show that VAChT is no longer correlated with ChAT or GABAergic markers. Yellow bars in (H) highlight the loss of VAChT correlations within cortical VIP terminals compared to (F).

**Figure S5.**
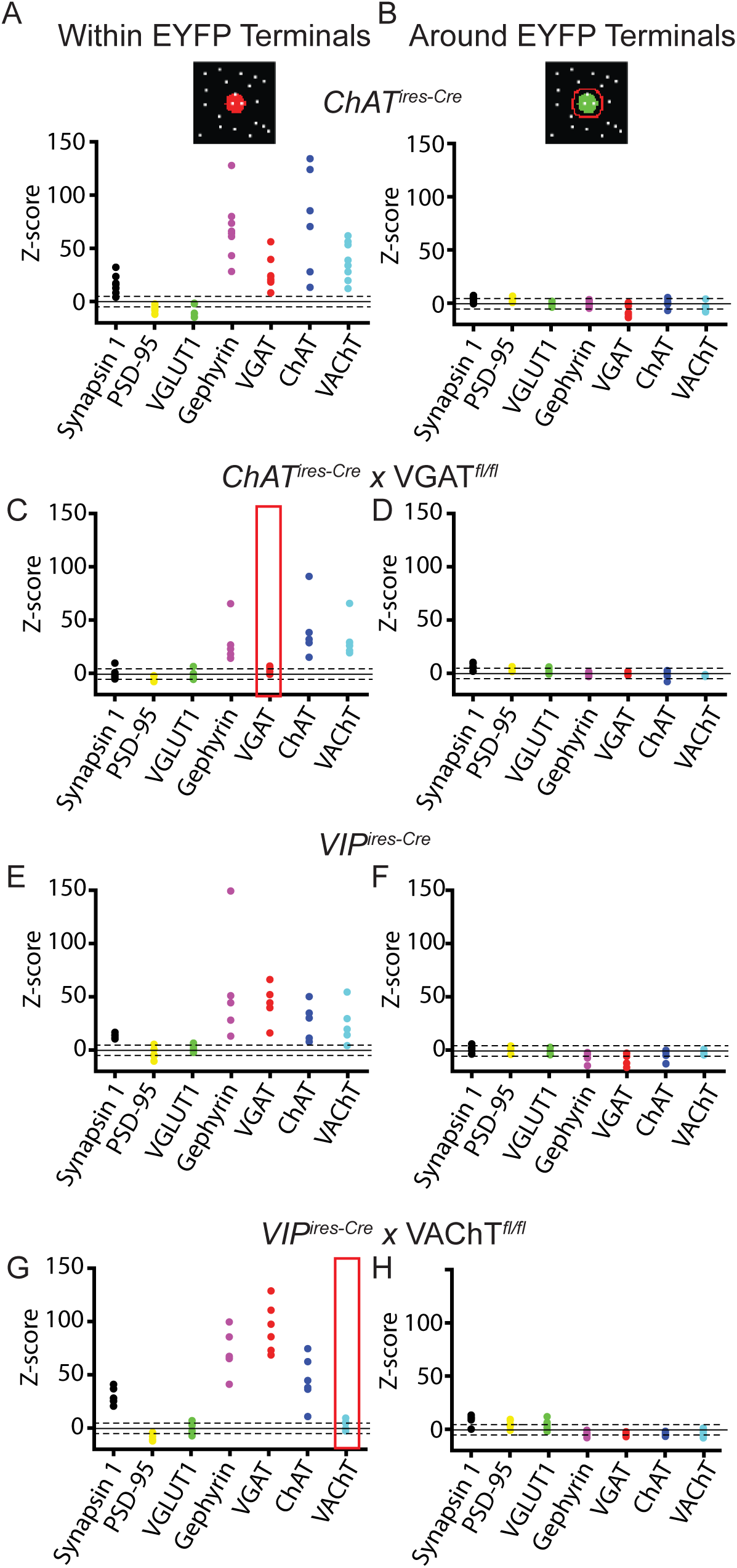
Summary antibody colocalization with VCIN and VIP pre-synaptic terminals. A,B) The z-scores of antibody colocalization with putative VCIN pre-synaptic terminals (A) and the area around VCIN terminals (B) from ChAT^ires-Cre^ mice injected in the frontal cortex with AAV(8)-DIO-Synaptophysin-YFP (n = 8 samples from 3 mice). C,D) The Z-scores of antibody colocalization with VCIN pre-synaptic terminals (A) and the area around VCIN terminals (B) from mice with VGAT deleted from cholinergic neurons (ChAT^ires-Cre^ x VGAT^fl/fl^; n = 6 from 3 mice). Loss of VGAT enrichment is highlighted in red. E,F) Z-scores of antibody colocalization with VIP pre-synaptic terminals from VIPires-Cre mice injected with AAV(8)-DIO-Synaptophysin-EFYP (n = 5 from 3 mice). G,H) Z-scores of antibody colocalization with VIP pre-synaptic terminals from mice with VAChT deleted from VIP neurons (VIP-^ires-Cre^ x VAChT^fl/fl^; n = 6 from 3 mice). Loss of VAChT enrichment is highlighted in red. Portions of this figure are duplicated from Figure 3.

**Figure S6.**
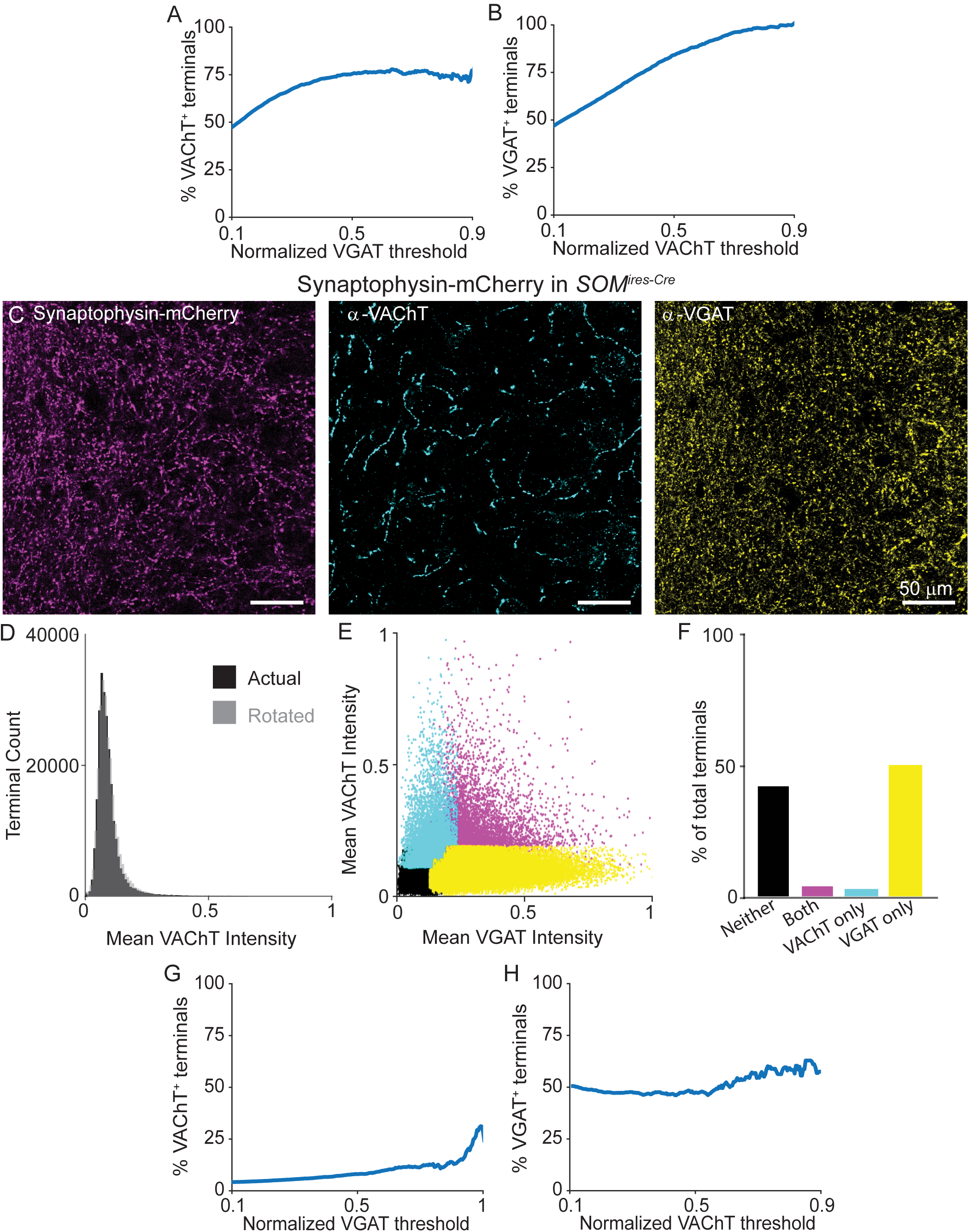
VAChT is present in a subset of terminals across intensity thresholds, and is absent from the terminals of Sst interneurons. A) The percent of terminals that are positive for VAChT (as determined by a mean intensity threshold greater than the Otsu threshold) across a range of VGAT intensity thresholds. As the threshold for VGAT intensity increases, the percent of VAChT+ terminals plateaus around 70%. B) The percent of terminals that are positive for VGAT (also determined by a mean intensity threshold greater than the Otsu threshold) across a range of VAChT intensity thresholds. The percent of VGAT+ terminals increases to 100% as the VAChT intensity threshold increases. This indicates that there are two main populations of terminals – those that express both VGAT and VAChT, and those that express only VGAT. C) Example images of Sst+ pre-synaptic terminals labeled with AAV(8)-DIO-Synaptophysin-mCherry injected into the cortex of SOM^ires-Cre^ mice. Left: Synaptophysin-mCherry; Middle: VAChT immunostain; Right: VGAT immunostain. D) Histogram of mean VAChT fluorescence intensity within Synaptophysin-mCherry+ pre-synaptic terminals of Sst interneurons. Black histogram represents the actual VAChT intensities, the grey histogram represents the mean VAChT intensities when the mCherry+ mask is rotated 90° relative to the VAChT immunostain. E) Scatter plot of mean VGAT and VAChT intensity in each putative mCherry+ pre-synaptic terminal (n = 249,899 putative terminals from 14 image stacks from 2 mice). Terminals are color-coded according to expression of VAChT and VGAT (Black – neither VGAT or VAChT, Magenta – both VGAT and VAChT, Cyan – VAChT only, Yellow – VGAT only). F) Quantification of the number of terminals of each type in (E). G,H) The percent of VAChT+ terminals (G) and VGAT+ terminals (H) across a range of VGAT and VAChT intensity thresholds, respectively. As VGAT intensity threshold increases, there is no relationship with the percent of terminals that are VAChT+ (G). Likewise, as VAChT intensity threshold increases, there is no relationship with the percent of terminals that are VGAT+ (H).

**Figure S7.**
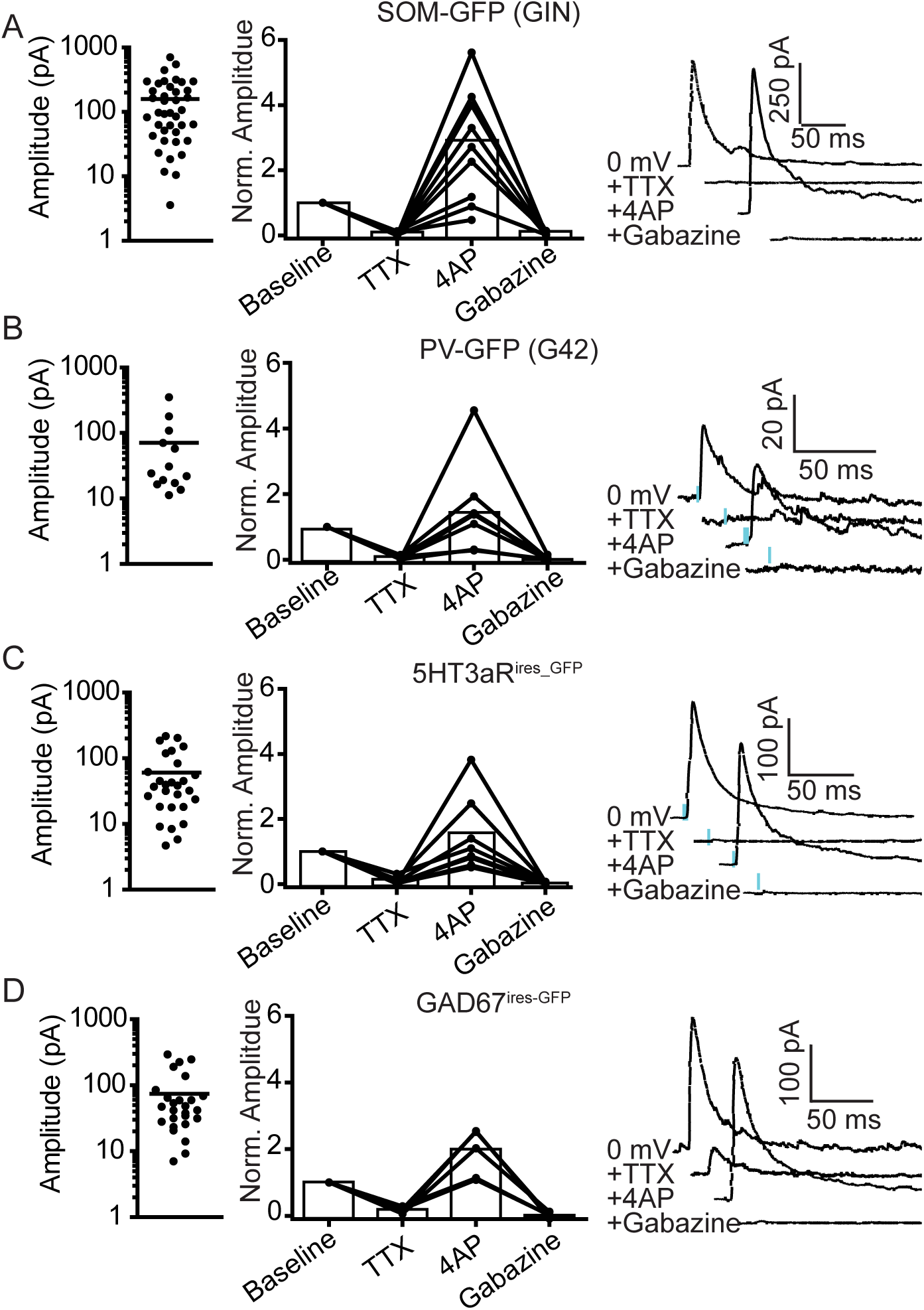
GABA_A_R-mediated synaptic currents from VCINs are mono-synaptic. (A-D). Synaptic response amplitudes and sensitivity to TTX, 4AP, and gabazine for GFP-expressing post-synaptic neurons in SOM-GFP (GIN) mice (A), PV-GFP (G42) mice (B), 5HT3aR^ires-GFP^ mice (C), and GAD67^ires-GFP^ mice. Left: mean response amplitude. Middle: Quantification of sensitive to TTX, 4AP, and gabazine. All synaptic responses are sensitive to TTX, substantially rescued by 4AP, and completely abolished by gabazine. Right: example traces showing synaptic response amplitude following consecutive application of TTX, 4AP, and gabazine.

**Figure S8.**
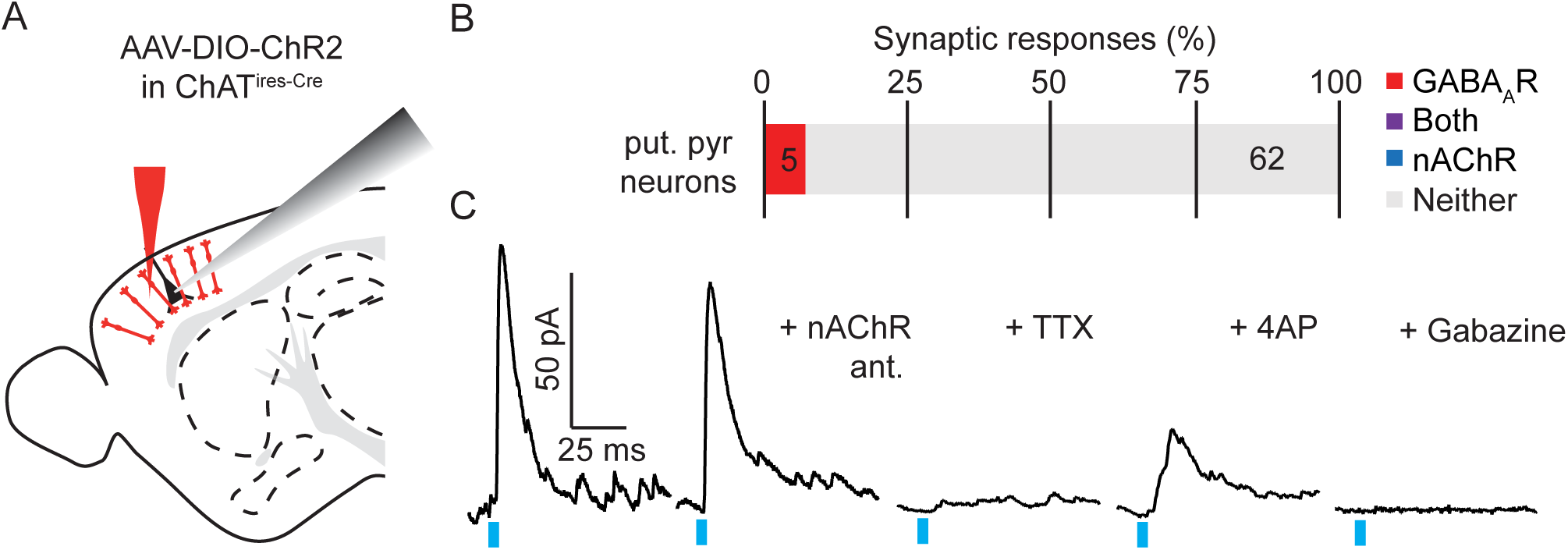
VCINs show low rate of connectivity to putative pyramidal neurons. A) Experimental design: AAV(8)-DIO-ChR2 was injected into the frontal cortex of ChATires-Cre mice. Following 3-4 weeks to allow for virus expression, whole-cell voltage clamp recordings of pyramidal neurons were made from acute sagittal slices. Pyramidal neurons were identified based on their morphology and laminar position. B) The proportion of pyramidal neurons with synaptic responses to optogenetic stimulation of VCINs. The number of cells per category are indicated. C) Example GABAA receptor-mediated synaptic current, which is insensitive to nAChR antagonists DHβE, MLE, and MEC, is blocked by TTX and subsequently rescued by 4AP, and completely blocked by gabazine.

**Figure S9.**
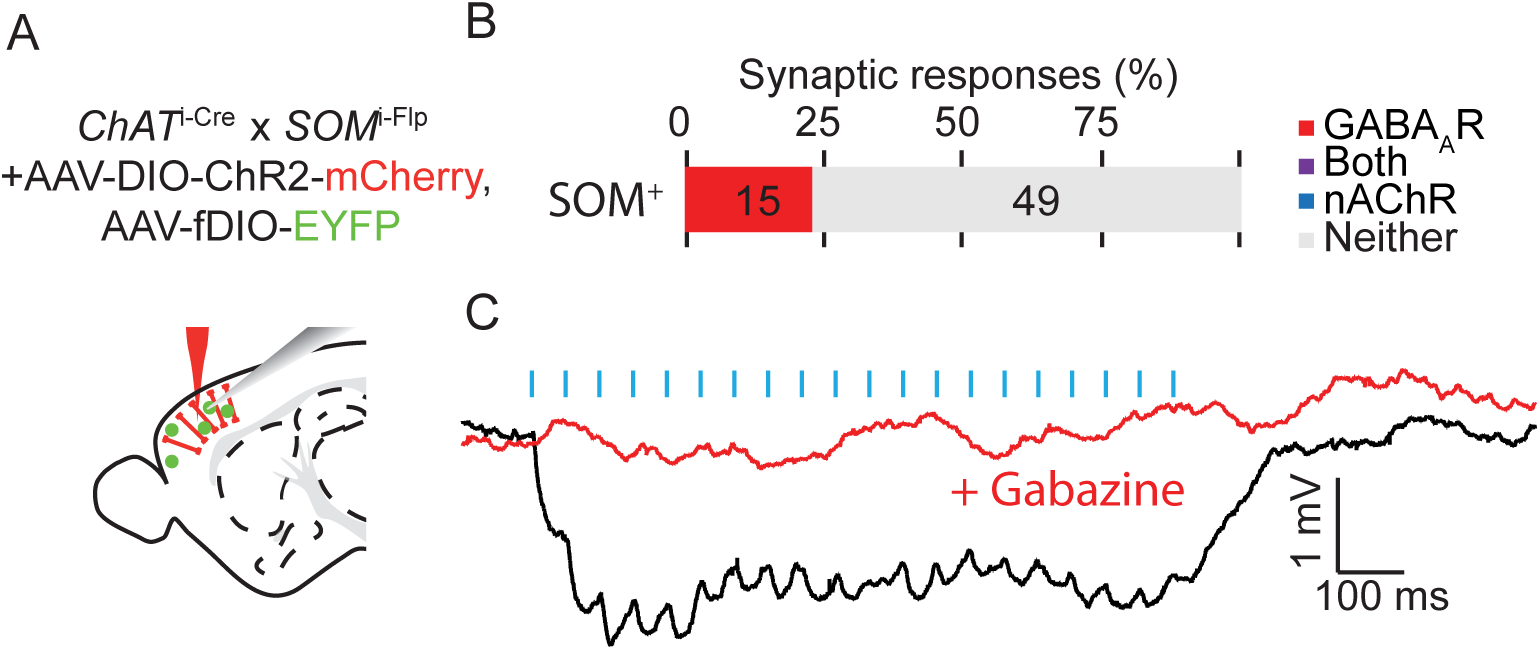
VCINs inhibit a subset of deep layer Sst+ interneurons. (A) Experimental design: Because the SOM-GFP (GIN) BAC transgenic mouse line does not express in deeper layer Sst+ interneurons, we targeted Sst+ interneurons in Layer 5 and Layer 6 by crossing ChAT^ires-Cre^ mice with SOM^ires-flp^ mice and injecting with a AAV(8)-DIO-ChR2-mCherry and AAV(DJ)-fDIO-EYFP. Whole-cell current clamp recordings were obtained from EYFP+ neurons in layers 5 and 6 after 3-4 weeks of virus expression from acute sagittal slices. (B) The proportion of cells showing synaptic responses following optogenetic stimulation of VCINs. GABAergic responses were identified based on hyperpolarizing post-synaptic potentials and sensitivity to gabazine. The number of neurons in each category are indicated. (C) Example synaptic response to a train of optogenetic stimulation (20 x 20 Hz, 3 ms pulses) showing hyperpolarization that is blocked by gabazine.

**Figure S10.**
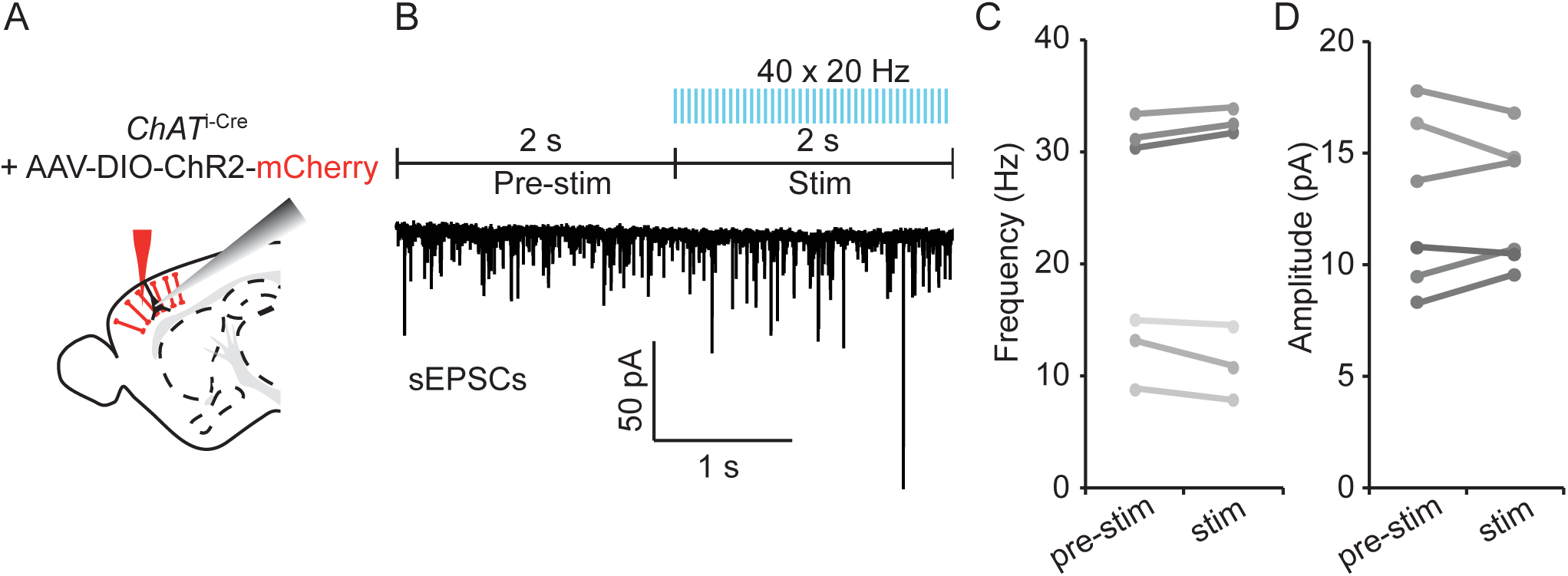
VCINs have no effect on pre-synaptic release properties. A,B) Experimental Paradigm: AAV(8)-DIO-ChR2-mCherry was injected into the frontal cortex of ChAT^ires-Cre^ mice and acute sagittal slices cut after 3-4 weeks. Spontaneous excitatory post-synaptic currents (sEPSCs) were recorded during whole-cell voltage clamp of Layer 2/3 pyramidal neurons. Several seconds of baseline sEPCSs were gathered, followed by 2 seconds of optogenetic stimulation of ChR2+ neurons (40 x 20 Hz, 3 ms pulses). (C) sEPCS frequency (left) and amplitude (right) before and during stimulation.

**Figure S11.**
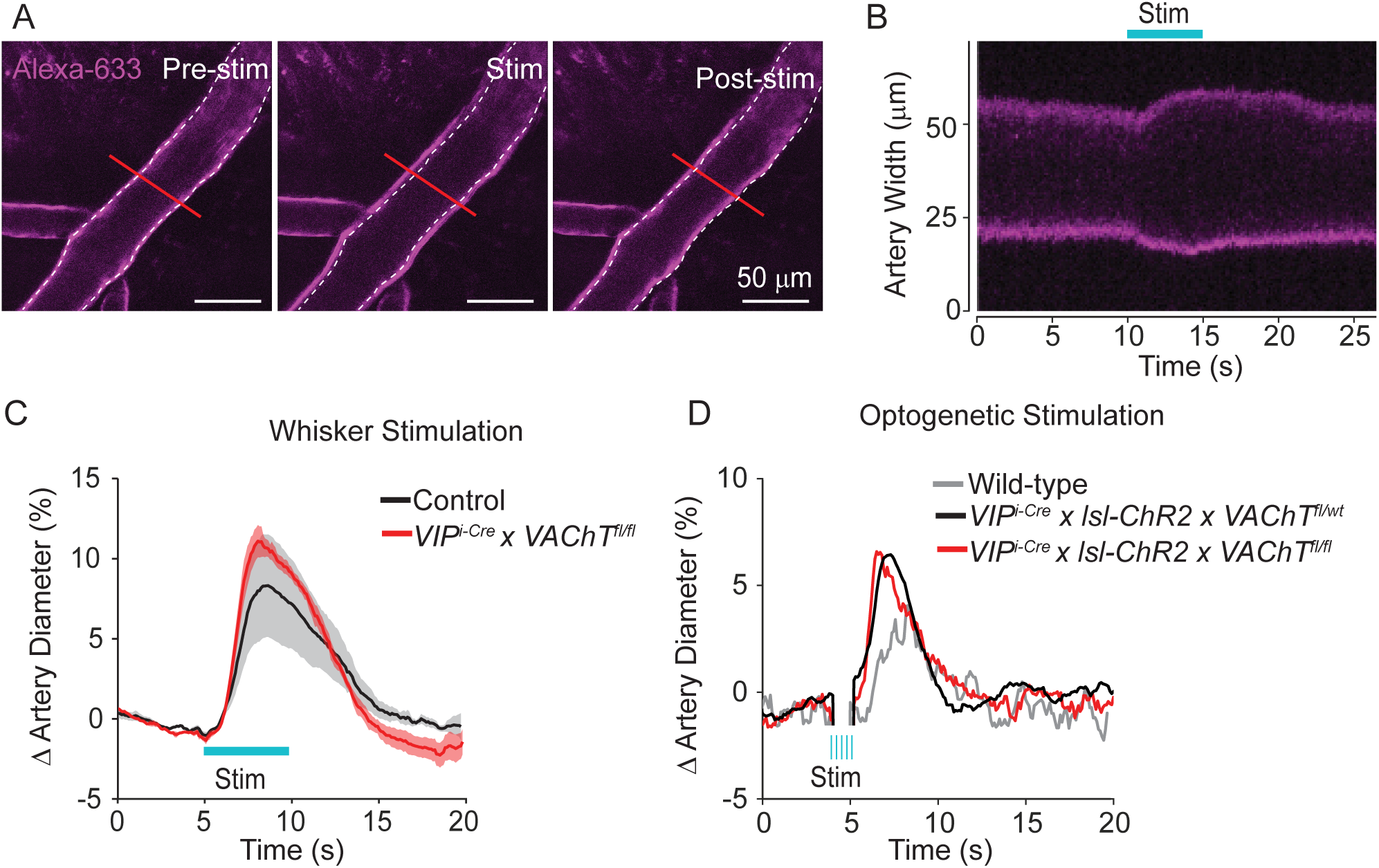
ACh release from VIP+ interneurons is not necessary for neurovascular coupling. A) Example images of pial arteries imaged in vivo over barrel cortex following retro-orbital injection of Alexa 633 hydrazide both prior to (left), immediately following (middle), and several seconds following (right) 3 second stimulation of the contralateral whiskers. B) Kymograph of the red line illustrated in (A) showing the width of the artery over time. C) Average percent change in pial artery diameter imaged over the barrel cortex of from 3 mice lacking VAChT in VIP+ interneurons (VIP^ires-Cre^ x VAChT^fl/fl^) and 3 heterozygous or wild-type littermate controls during contralateral stimulation of whiskers. D) Average percent change in pial artery diameter imaged over the barrel cortex during optogenetic stimulation (5 x 20 hz, 5 ms pulses) from 3 mice lacking VAChT in VIP+ interneurons (VIP^ires-Cre^ x Rosa26^lsl-ChR2-EYFP^ x VAChT^fl/fl^), 2 heterozygous littermate controls (VIP^ires-Cre^ x Rosa26^lsl-ChR2-EYFP^ x VAChT^fl/wt^), and 2 wild-type controls that do not express ChR2.

